# Transcriptomic entropy benchmarks stem cell-derived cardiomyocyte maturation against endogenous tissue at single cell level

**DOI:** 10.1101/2020.04.02.022632

**Authors:** Suraj Kannan, Michael Farid, Brian L. Lin, Matthew Miyamoto, Chulan Kwon

## Abstract

The immaturity of pluripotent stem cell (PSC)-derived tissues has emerged as a universal problem for their biomedical applications. While efforts have been made to generate adult-like cells from PSCs, direct benchmarking of PSC-derived tissues against *in vivo* development has not been established. Thus, maturation status is often assessed on an *ad-hoc* basis. Single cell RNA-sequencing (scRNA-seq) offers a promising solution, though cross-study comparison is limited by dataset-specific batch effects. Here, we developed a novel approach to quantify PSC-derived cardiomyocyte (CM) maturation through transcriptomic entropy. Transcriptomic entropy is robust across datasets regardless of differences in isolation protocols, library preparation, and other potential batch effects. With this new model, we analyzed over 45 scRNA-seq datasets and over 52,000 CMs, and established a cross-study, cross-species CM maturation reference. This reference enabled us to directly compare PSC-CMs with the *in vivo* developmental trajectory and thereby to quantify PSC-CM maturation status. We further found that our entropy-based approach can be used for other cell types, including pancreatic beta cells and hepatocytes. Our study presents a biologically relevant and interpretable metric for quantifying PSC-derived tissue maturation, and is extensible to numerous tissue engineering contexts.

**Significance Statement:** There is significant interest in generating mature cardiomyocytes from pluripotent stem cells. However, there are currently few effective metrics to quantify the maturation status of a single cardiomyocyte. We developed a new metric for measuring cardiomyocyte maturation using single cell RNA-sequencing data. This metric, called entropy score, uses the gene distribution to estimate maturation at the single cell level. Entropy score enables comparing pluripotent stem cell-derived cardiomyocytes directly against endogenously-isolated cardiomyocytes. Thus, entropy score can better assist in development of approaches to improve the maturation of pluripotent stem cell-derived cardiomyocytes.

---

**T**he development of robust protocols for differentiation of pluripotent stem cells (PSCs) to a range of somatic tissues has represented a huge advance in biomedical research over the past two decades. Many somatic cell types are non-proliferative and difficult to obtain from patients, and thus PSCs may be the most viable source for generating large quantities of specialized tissue. PSC-derived tissues have numerous promised applications in regenerative medicine, drug efficacy and toxicity screening, and *in vitro* disease modeling (1–5). However, clinical application of PSC-derived tissues has been limited thus far due to the failure of these cells to mature to a fully adult-like phenotype *ex vivo*. This phenomenon has been observed in a range of engineered tissue types, including cardiomyocytes (CMs) (6), hepatocytes (7), pancreatic islet cells (8), neurons (9), and others, and represents a major biomedical hurdle.

To date, numerous engineering approaches have been proposed to improve PSC-derived tissue maturation. These approaches have included cocktails, induction of physical stimuli, co-culture with other cells, and construction of three-dimensional tissues, typically with the goal of recapitulating the native milieu (4, 6, 8, 10–14). Benchmarking the efficacy of these interventions has been challenging, however, as functional assays for direct comparison of PSC-derived cells to endogenous adult cells are often technically infeasible. Thus, most engineered tissues are compared either to a two-dimensional *in vitro* control or at best one discrete (usually neonatal) *in vivo* timepoint, rather than across the continuous spectrum of *in vivo* maturation. Several groups have proposed use of -omics data to compare engineered to *in vivo* tissues for certain cells (15–18). However, these approaches have been limited to bulk samples, which precludes their use when PSC differentiation yields highly heterogeneous populations.

scRNA-seq has emerged as a powerful tool for measuring the transcriptomes of large numbers of single cells, and is an intriguing candidate for new metrics of tissue maturation. Unfortunately, differences in isolation protocols, library preparation methods, and sequencing machines, among other factors, can imbue scRNA-seq data with batch effects that are difficult to deconvolve (19, 20). In turn, this makes it difficult to directly compare expression of individual genes across datasets. While batch correction algorithms have been developed (21), they are primarily designed for correcting or integrating datasets with multiple well-defined cell types with significantly different gene expression patterns rather than one continuously evolving cell type. Thus, an optimal scRNA-seq-based metric of maturation must facilitate direct comparison of maturation status while being robust to batch effects.

Here, we developed an approach based on quantifying gene *distributions* to assess PSC-derived tissue maturation. Given the significant burden of cardiac disease (22), we focused our analysis on CMs, the primary contractile cells of the heart. Our approach is based on the generally-observed phenomenon that less differentiated cells are typically more promiscuous in their transcriptional activities of signaling pathways, leading to a diverse gene expression profile. However, as they differentiate, they prune unnecessary signaling pathways and hone in on a relatively narrow gene expression profile (23). This observation has been leveraged in several previous approaches to study differentiation of stem cells to progenitors and subsequently to committed lineages (24–29). We applied this principle to study the maturation of committed CMs by developing a metric based on a modification of the Shannon entropy of scRNA-seq gene expression data. Our transcriptomic entropy-based metric not only adequately stages single CMs, but that scores are consistent across datasets regardless of potentially confounding batch effects. Using datasets from the literature, we performed a meta-analysis of CM maturation based on transcriptomic entropy. We subsequently demonstrated the use of our approach to infer the maturation status of PSC-CMs. While our primary focus was on CMs, we also showed initial evidence of applicability to other celltypes, in particular pancreatic beta cells and hepatocytes. These results establish transcriptomic entropy as a viable metric for benchmarking PSC-derived tissue maturation.

## Results

### scRNA-seq CM reference captures maturation-related changes

As a first step, we sought to construct a reference scRNA-seq library for CM maturation. Sequencing of postnatal CMs, which are large and fragile, has been previously limited (30). Recently, however, we developed a method to isolate healthy adult CMs to generate high quality scRNA-seq libraries using large-particle fluorescence-activated cell sorting (LP-FACS) (31). We used this approach to isolate CMs from Myh6-Cre; mTmG (*α*MHC x mTmG) mice, in which cells expressing cardiac-specific myosin heavy chain are readily separated by GFP expression (**Figure 1a**). We generated a library of ∼1000 CMs from 12 points over the course of maturation. Our reference particularly sampled cells within the first three weeks postnatally, as this period may be critically relevant to the maturation process but is underrepresented in existing CM scRNA-seq datasets.

**Fig. 1.**
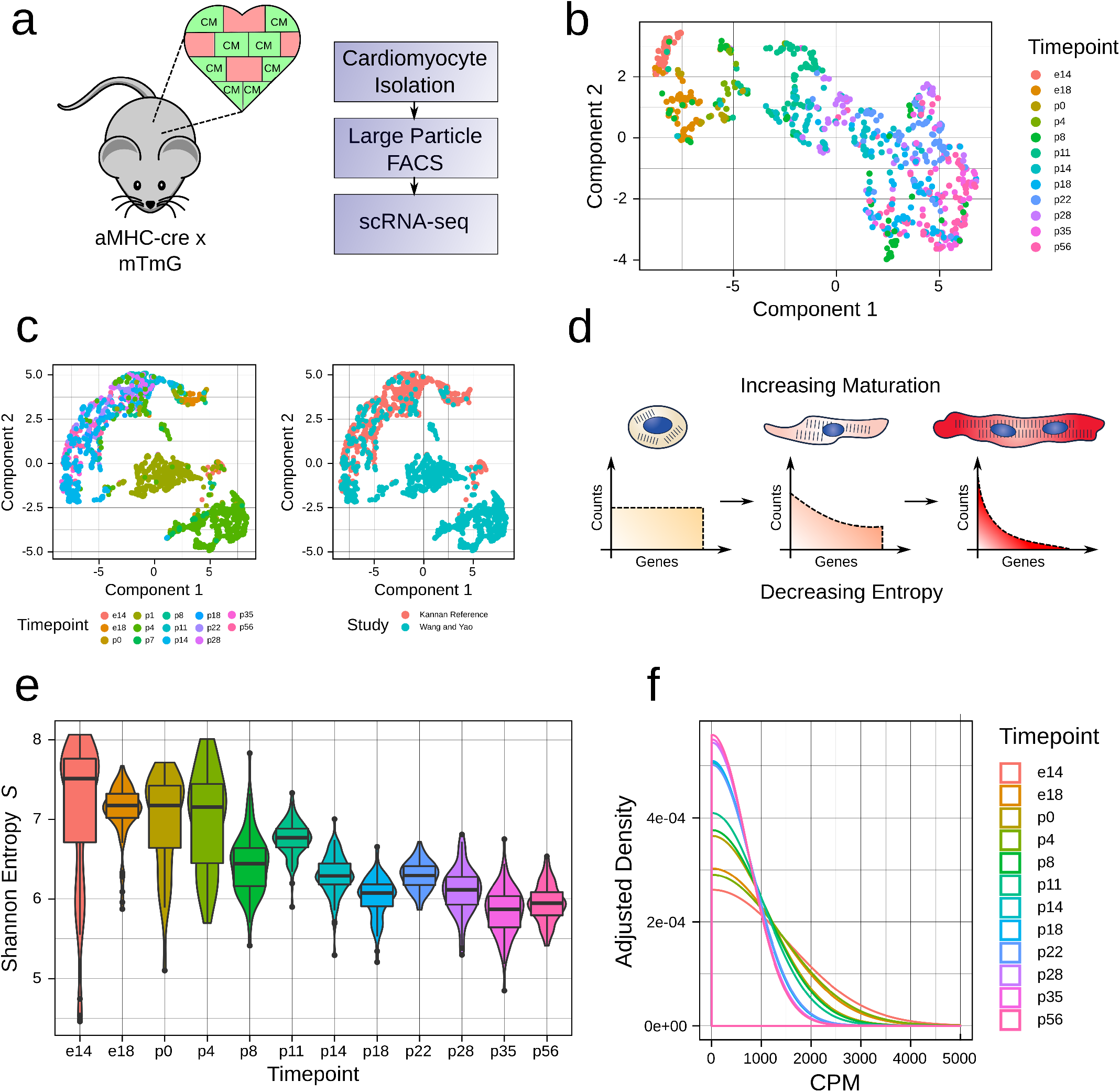
scRNA-seq constructs a reference for CM maturation. **a**. Mouse model used to generate perinatal maturation reference scRNA-seq library. In the *α*MHC-cre x mTmG mouse, CMs are labeled by GFP. **b**. UMAP dimensionality reduction (via Monocle 3) for the maturation reference. **c**. mnnCorrect-based integration of Wang and Yao et al. dataset with reference dataset. **d**. Our model for changes in gene distribution over CM maturation. As CMs undergo the maturation process, they transition from a broad gene distribution (characterised by high entropy) to a more narrow distribution (characterised by low entropy). **e**. Shannon Entropy *S* computed for each timepoint in the maturation reference dataset. **f**. Smoothed density estimates for genes expressed at 0-5000 counts per million (CPM) for each timepoint in the maturation reference dataset.

Dimensionality reduction of our reference with uniform manifold approximation and projection (UMAP) revealed a continuum of maturation from e14 in p56, in concordance with previous results from our group and others (32–34) (**Figure 1b**). As an initial strategy, we considered quantification of maturation status by integration of query datasets with our reference. We tested this approach by using mnnCorrect (35) to combine other *in vivo* CM scRNA-seq datasets with our reference while correcting study-specific batch effects. While this approach worked for some datasets, in other cases study-dependent batch-effects were only partially corrected (**Figure 1c**). Moreover, integration often changed cell-to-cell distances within our original reference itself. These results confirmed the difficulty of quantifying CM maturation through a conventional dimensionality reduction/batch correction-based pipeline, and prompted us to seek other approaches for quantifying maturation from scRNA-seq data.

### Shannon entropy of single cell gene expression decreases over CM maturation

We next considered an approach based on gene distribution changes. Following terminal differentiation, CMs undergo a lengthy maturation process characterized by gradual and unidirectional changes in gene expression (18). Based on previous findings, we proposed a model for transcriptional maturation of CMs analogous to cellular differentiation (**Figure 1d**). In this model, nascent cardiomyocytes express a broad gene expression profile. However, as they mature, they slowly reduce expression of immature gene pathways (e.g. cell cycle) while upregulating genes required for mature function (e.g. sarcomere, calcium handling, oxidative phosphorylation). These gradual changes in gene distribution can be quantified by established diversity metrics such as the well-known Shannon entropy. In our model, immature myocytes will present with high transcriptomic entropy, which subsequently decreases in a continuous manner over the course of maturation.

To test the validity of this model, we computed the Shannon entropy *S* on the unique molecular identifier (UMI) counts of our maturation reference (**Figure 1e**). Entropy gradually decreased from e14 to p56, with a notable shift from p4 to p8, thereby supporting our hypothesized entropy model. We additionally plotted the averaged gene distributions for each timepoint (**Figure 1f**). As expected, earlier timepoints showed a more broad distribution compared to later timepoints. These results supported the use of Shannon entropy to quantify CM maturation status from scRNA-seq data.

### Gene and cell filtration are necessary for cross-study comparison

Given the correspondence between Shannon entropy and CM maturation status, we next sought to determine whether we could extend our transcriptomic entropy model to many CM scRNA-seq datasets generated across multiple labs. We identified publicly available scRNA-seq datasets containing CMs isolated *in vivo* (**Supplementary Table 1**). Our meta-analysis included 34 mouse datasets and 5 human datasets spanning numerous timepoints across the range of development. Additionally, the collected datasets represented significant diversity in terms of isolation methods, sequencing protocols, mapping/counting pipelines, and datatypes (including reads from full-length scRNAseq protocols, 3’ counts from UMI protocols prior to UMI collapsing, and UMIs). However, several technical challenges prevented accurate cross-study comparison of Shannon entropy computed on raw, unfiltered datasets. These particular challenges and our solutions are addressed here.

#### Handling multiple mapping/counting pipelines

One problem that was observed in certain sequencing mapping/counting pipelines was the incorrect mismapping of mitochondrial reads to pseudogenes. In mice, fragments of the mitochondrial genome are present as pseudogenes in the nuclear genome (termed nuclear mitochondrial insertion sequences (36)). These fragments often show identical or near-identical sequences to mitochondrial genes. Thus reads are often multi-mapped between canonical mitochondrial genes and pseudogenes, leading to inaccurate gene quantification in pipelines counting multi-mapping reads. This issue was particularly problematic for CMs, as they naturally express high amounts of mitochondrial genes (31).

As our goal was to enable entropy to be widely usable across many protocols, we included an approximate pseudogene correction in our pipeline. We identified cross-mappings between pseudogenes and canonical genes, and subsequently removed all pseudogene counts and added them to the corresponding canonical mitochondrial genes. To test the efficacy of our pseudogene correction, we tested the entropy score and well as mitochondrial gene percentages before and after correction for several mapping/counting methods (**Figure 2a, Supplementary Figure 1**). As a genomic method, we used the zUMIs pipeline (37), which uses STAR for mapping followed by FeatureCounts for counting. In this method, multimapping counts are effectively randomly allocated between mitochondrial reads and pseudogenes. As a transcriptomic method, we utilized kallisto|bustools (38). We used kallisto|bustools with two indices - a full index containing all mouse cDNAs from ENSEMBL (kb.full), and an index containing only protein coding, lincRNAs, and antisense RNAs analogous to the Cell Ranger index (kb.cellranger). Lastly, we also used Cell Ranger, a part of the 10x Genomics pipeline. The Cell Ranger index does not contain pseudogenes, and thus does not feature mitochondrial read mismapping.

**Fig. 2.**
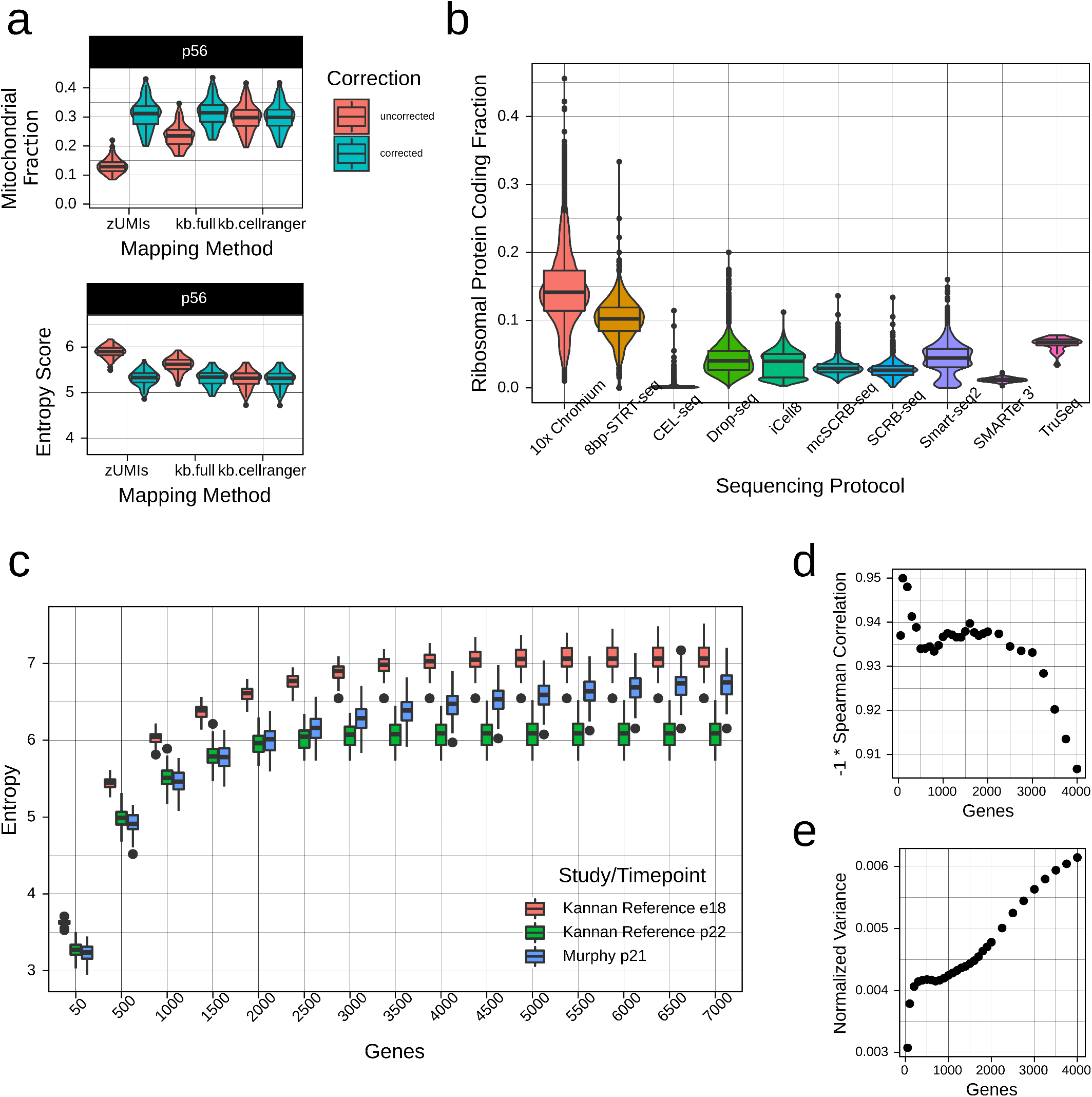
Gene filtration facilitates cross-study comparisons of entropy. **a**. Correction of mismapped mitochondrial reads. The use of a genomic mapping algorithm (such as zUMIs) or a “full” reference containing pseudogenes can lead to erroneous mapping of mitochondrial genes to pseudogenes, in turn inflating entropy score. We included a correction step that facilitates usage of data from a range of mapping pipelines. **b**. Proportion of ribosomal protein coding genes in mouse *in vivo* datasets, grouped by library preparation method. Given the clear protocol-dependence of these genes, we eliminated them from analysis. **c**. Entropy at different gene subsamplings for two studies with different sensitivities. Data from e18 and p22 from the maturation reference and p21 from Murphy et al. are shown. **d**. Spearman correlation between entropy and timepoint for different gene subsamplings (using median of entropy for each timepoint and study). **e**. Normalized variance of entropy for different gene subsamplings (using median of entropy for each timepoint). We normalized by scaling entropy at every subsampling to [0, 1].

The results of correction are shown for p56 samples in our reference (**Figure 2a**). Prior to correction, zUMIs and kb.full produced datasets with lower mitochondrial read percentage and therefore higher entropy. However, post-correction, these datasets showed entropy and mitochondrial read percentages that were nearly identical to kb.cellranger and Cell Ranger. Thus, datasets prepared using methods that include multi-mapping reads will be sufficiently corrected for cross-study comparison.

#### Ribosomal protein-coding genes

By default, we included for analysis genes with gene biotype “protein coding,” “antisense,” or “lncRNAs,” so as to focus on the key players of the transcriptome. We additionally considered ribosomal protein-coding genes, and found significant protocol-related biases in terms of expression of these genes (**Figure 2b, Supplementary Figure 2**). In particular, 10x Chromium and STRT-seq datasets appeared to have systematically higher percentages of ribosomal protein-coding genes than other protocols. This observation anecdotally matches observations made by others and likely indicates a protocol bias, though we are unsure about the reason this occurs. Therefore, we removed all ribosomal protein-coding genes prior to computation of entropy.

#### Variations in study sensitivity

Different scRNA-seq datasets will invariably detect different numbers of genes as a consequence of differences in sequencing depth and sequencing protocol sensitivity (39). However, higher sensitivity can lead to artificially higher entropy simply by inclusion of more terms in the summation. An example of this effect is shown in **Figure 2c**, where we compared our reference (median 3000 genes/cell) against our previously generated dataset (Murphy et al., median 7044 genes/cell). We thus explored subsampling of genes as a way to standardize for the summation term in entropy and therefore sensitivity differences. Selecting the optimal number of genes for subsampling required balancing two priorities. Too many genes would result in overemphasis on sensitivity differences and incorrect separation of cells with similar developmental stage, as above. However, as entropy does not scale linearly with subsampled genes, too few genes would result in compression of the dynamic range of the metric and similar scores for cells at different developmental stages. We optimized the former by computing by seeking to maximize Spearman correlation between entropy and timepoint (**Figure 2d**), and the latter by seeking to maximize the variance of normalized entropy across timepoints at each subsampling (**Figure 2e**). We selected 1000 genes as a reasonable subsampling based on both sets of results, though we would have obtained comparable results for 400-1500 genes.

#### Identifying poor quality datasets

Not all of the datasets identified were of sufficient quality for downstream analysis. This issue is particularly severe for CMs, as adult CMs are highly difficult to isolate at the single cell level by a number of classical methods, such as conventional FACS, single cell picking, or microfluidic devices such as the Fluidigm C1 (31). To identify such datasets, we used the percentage of mitochondrial reads as a quality control metric, discarding datasets with unusually high percentages (**Figure 3a**). Currently, there is no automated approach for easily identifying poor quality datasets. We thus erred on the side of caution, and tried to avoid eliminating datasets without clear rationale for doing so. We outlined our rationale for discarding any datasets in the **Appendix**, with the hope that transparency could suffice in the current absence of more rigorous disqualification criteria.

**Fig. 3.**
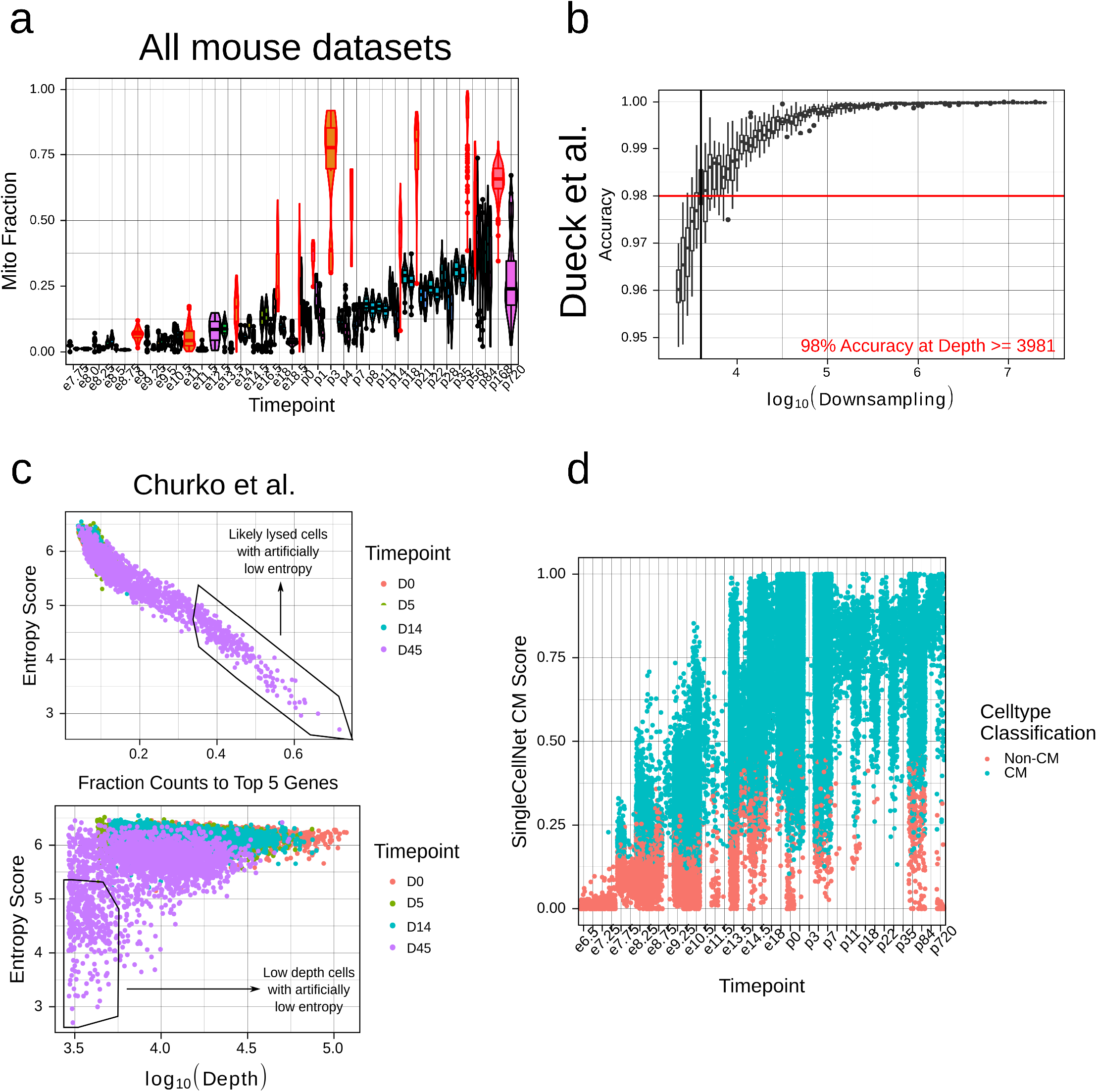
Standardized cell and study filtration enables meta-analysis of CM maturation with entropy. **a**. Mitochondrial gene fractions in mouse *in vivo* datasets. Datasets with unusually high proportions are highlighted in red and were removed from subsequent analysis. **b**. Subsampling of count depth in Dueck et al. dataset (the highest depth dataset in our analysis). We subsampled to a depth where the median number of genes remained > 1000. We subsequently computed the accuracy as deviation from baseline entropy. **c**. Unusually low entropy cells due to high top 5 gene percentage (top) or low depth (bottom) in the Churko et al. dataset. **d**. SingleCellNet CM scores for mouse *in vivo* datasets by timepoint. Cells are labeled based on whether their highest classification was for “cardiac muscle.”

In addition, we assessed the maximum dataset depth necessary for a study to be appropriately quantified by entropy. We tested several datasets with a range of baseline depths, and performed subsampling to determine a minimum required depth for accurate entropy quantifications (**Figure 3b, Supplemental Figure 3**). We defined accuracy based on the deviation from the baseline entropy, and set a threshold of 98% accuracy (corresponding to ∼0.1 change in entropy). Entropy was relatively robust to subsampling, with 98% accuracy being achieved at above ∼2000-4500 counts/cell, depending on the dataset. While this depth was sufficient for most of our assayed datasets, some very low-depth datasets were affected - in particular, all four Drop-seq datasets tested had depths ranging from 1500 - 4100 counts/cell. Given these results, we omitted the Drop-seq datasets from further analysis.

#### Quality control of poor-quality cells

Outside of dataset filtering, within-dataset quality control is an essential step in all scRNA-seq protocols (40–42). Protocols will inevitably generate cells that have been lysed or damaged, making them unsuitable for downstream analysis. As our study involves a meta-analysis of many independently generated datasets, we aimed to establish a standardized approach for quality control. This had the dual benefits of ensuring at least a minimal level of comparability while limiting the need to determine individual thresholds for each dataset. We focused on two primary metrics of quality control - cell depth and percentage of reads going to the top 5 highest expressed genes in each cell. We selected these metrics because we observed that they most affected quantification of entropy score (**Figure 3c**). We then defined normalized metrics based on both measurements by dividing the respective measurement by the median of that measure in that study and in that timepoint. Thus, while comparable cross-study, the metrics could be considered with respect to potential biological and technical variation. We then set a standardized threshold across all studies (**Supplementary Figure 4**).

#### Identifying CMs

In terms of cell-type filtration, our input datasets were fairly heterogeneous, with some including only CMs while others were more broad. Thus, we used SingleCellNet (43) to identify and retain only cells with CM signature. SingleCellNet uses top-scoring pair to enable cross-platform comparisons of test data against a training dataset to annotate celltypes, and has performed well in benchmarking (44). We used the Tabula Muris (45) as a reference dataset to test against many celltypes. However, as the Tabula Muris is constructed on adult tissues, we were concerned that early-stage CMs may be poorly classified. We thus tested the predicted cell annotations from SingleCellNet across our mouse *in vivo* datasets. We classified a cell as a CM if its score for “cardiac muscle cell” was higher than the score for any other celltype. We found that, while prediction scores for CMs increased over time, CMs were identified as early as e7.5, corresponding appropriately to the onset of cardiomyogenesis (**Figure 3d**). In human *in vivo* datasets, CMs were present by embryonic week 5, which was the earliest timepoint for which we had data (**Supplemental Figure 5**). These results supported the use of the Tabula Muris reference with SingleCellNet, even for identifying nascent CMs.

### Entropy score enables cross-study inference of maturation status

Based on the previous results, we developed a workflow for addressing major technical confounding variables to enabling cross-study comparisons (**Figure 4a**). The output of our workflow is the computed Shannon entropy on the filtered datasets, which we refer to as *entropy score* through the remainder of the manuscript. We tested the utility of entropy score on our previously identified mouse and human *in vivo* CM datasets, which after filtration composed 36,436 CMs. Entropy score gradually decreased over developmental time, as hypothesized by our model (**Figure 4b, Supplementary Figure 6**). Notably, despite the marked heterogeneity of dataset characteristics, entropy score was consistent at similar timepoints across multiple datasets. In particular, entropy score showed remarkable concordance between datasets featuring different datatypes. For example, using four UMI-based datasets generated by our group, we found that the ratio of entropy score computed prior to versus after UMI collapsing was 1.02 (**Supplementary Figure 7**).

**Fig. 4.**
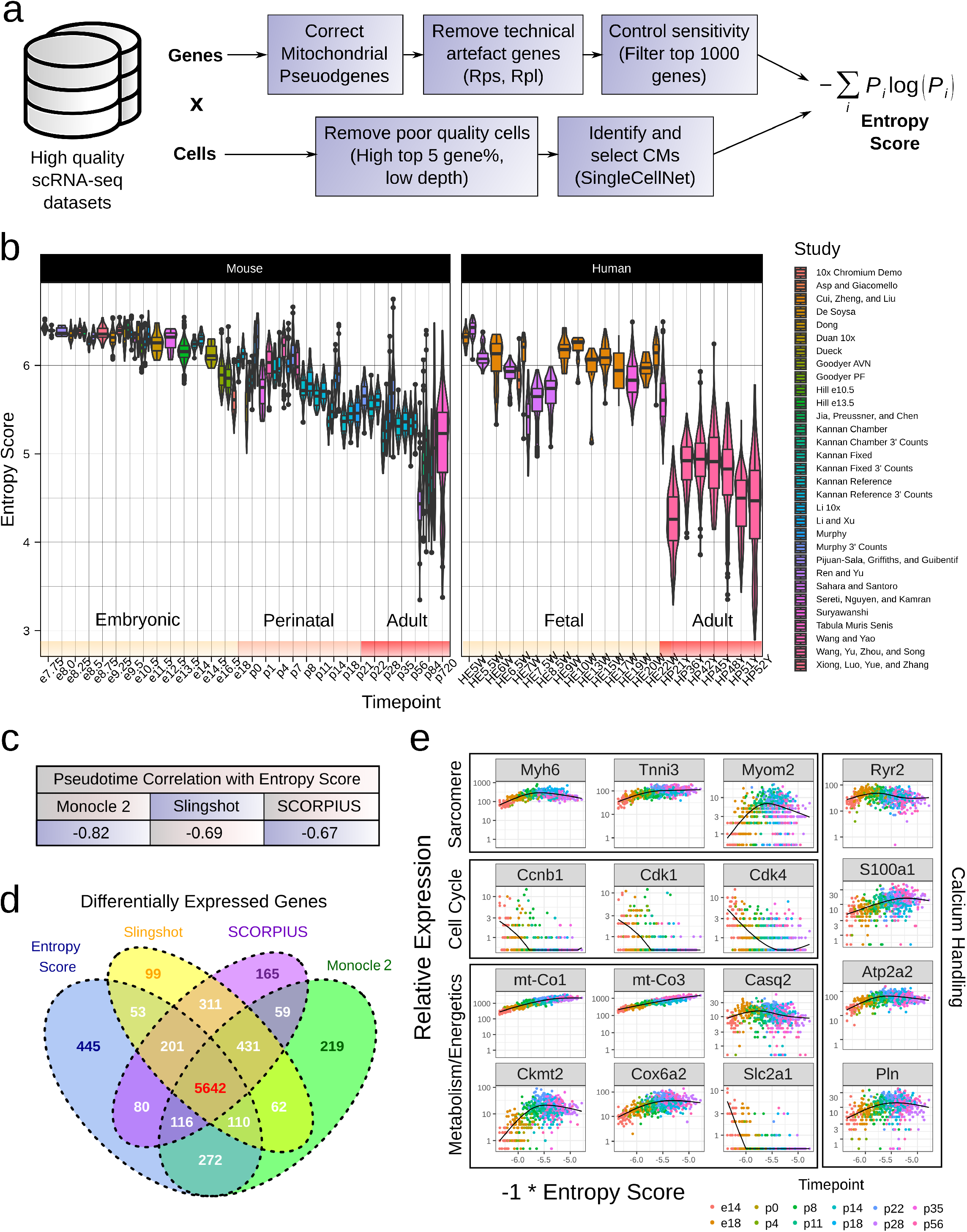
Entropy score enables cross-study and cross-species comparison of CM maturation status and recapitulates gene trends. **a**. Workflow for computing entropy score from high quality scRNA-seq datasets. **b**. Entropy score for mouse and human *in vivo* CMs taken from publicly available datasets. HEW = human embryonic week, HPY = human postnatal year. **c**. Pearson correlation between entropy score and calculated pseudotimes for our maturation reference dataset for three trajectory inference methods: Monocle 2, Slingshot, and SCORPIUS. **d**. Venn diagram showing overlap in identified differentially expressed genes between entropy score and trajectory inference methods. Differentially expressed genes were identified by fitting generalized additive models to gene trends over the corresponding pseudotime in Monocle 2, and selecting genes with adjusted p-value < 0.05. **e**. Gene expression trends over entropy score for genes involved in CM maturation, including sarcomeric, cell cycle, metabolism, and calcium handling genes.

In the mouse *in vivo* datasets, entropic changes occurred in three broad phases (**Figure 4b**). In the embryonic phase (∼e7.75-e16.5), entropy score decreased relatively slowly. Upon initiation of the perinatal phase at e16.5, entropy score decreased more rapidly before converging onto a mature adult-like phase at p21. These changes correspond well to previous literature about the dynamics of CM maturation, particularly regarding the perinatal CM maturation window (46).

We were additionally curious about the efficacy of entropy score to capture the maturation status of human CMs. We found that there was good concordance in entropy score between stage-matched mouse and human tissues (**Figure 4b**). In particular, fetal tissues (ranging from embryonic week 5 to embryonic week 22) corresponded approximately to e13.5-e14.5 in mice, while adult human CMs presented with entropy score comparable to adult mouse CMs. We did observe that one dataset (Sahara et al.) showed a notably lower entropy score at embryonic weeks 7-8, though we suspect this may have to do with dataset quality issues. Taken as a whole, however, these results support the use of entropy score as a cross-study, cross-species metric of CM maturation.

### Entropy score recapitulates gene expression trends in CM maturation

We next tested whether entropy score computationally ordered single CMs based on their progression along the maturation process, akin to so-called trajectory inference or pseudotime analysis methods. We selected three well-known trajectory inference methods - Monocle 2, Slingshot, and SCORPIUS - based on their performance in recent benchmarking studies, particularly with reconstructing unidirectional topologies (47). We then performed trajectory inference with our maturation reference dataset and compared the resultant pseudotimes with entropy score. Additionally, we identified genes differentially expressed over pseudotime/entropy score for each method respectively. Entropy score correlated only moderately with pseudotimes for the three methods (**Figure 4c, Supplementary Figure 8**). However, there was notable overlap in identified differentially expressed genes (**Figure 4d**). In particular, ∼93.6% of genes identified as differentially expressed over entropy score were also identified by at least one other method, and ∼81.5% were identified as differentially expressed by all methods. Moreover, when treated as a pseudotime metric, entropy score accurately recapitulated known CM maturation gene expression trends (**Figure 4e**). We further tested entropy score as a pseudotime metric in datasets composed of only one biological timepoint but a range of entropy scores. Intriguingly, gene expression trends across entropy score in these one-timepoint datasets largely matched the trends observed in our maturation reference dataset (**Supplementary Figure 9**). These results suggest that entropy score can effectively reconstruct the CM maturation trajectory as it occurs heterogeneously at the single cell level, and can accurate quantify single CM maturation status regardless of the biological timepoint of the sample.

### Human PSC-CMs do not mature beyond embryonic stage

Having validated entropy score as a metric of CM maturation *in vivo*, we next tested the entropy score of PSC-CMs from publicly available datasets (**Supplementary Table 2**). We identified 8 datasets of directed differentiation of human induced PSCs to CMs, and analyzed 13,171 cells between D(ay)9 and D100 of differentiation post-filtering. Though there was some variation from study to study (perhaps due to line-to-line differences or variations in differentiation protocol), there was modest decrease in entropy score over the course of differentiation (**Figure 5a**). However, no study generated CMs with entropy score lower than human fetal tissues, confirming the immature nature of PSC-CMs. Moreover, there was limited change in entropy score in PSC-CMs after D45, even with long-term culture up to D100, suggestive of maturation arrest. Interestingly, the entropy score of these later timepoint PSC-CMs corresponded to the initiation of the perinatal phase of mouse CM maturation *in vivo*. This observation may point to dysregulation of the endogenous perinatal maturation program during *in vitro* directed differentiation as a cause of poor PSC-CM maturation status, and merits further investigation.

**Fig. 5.**
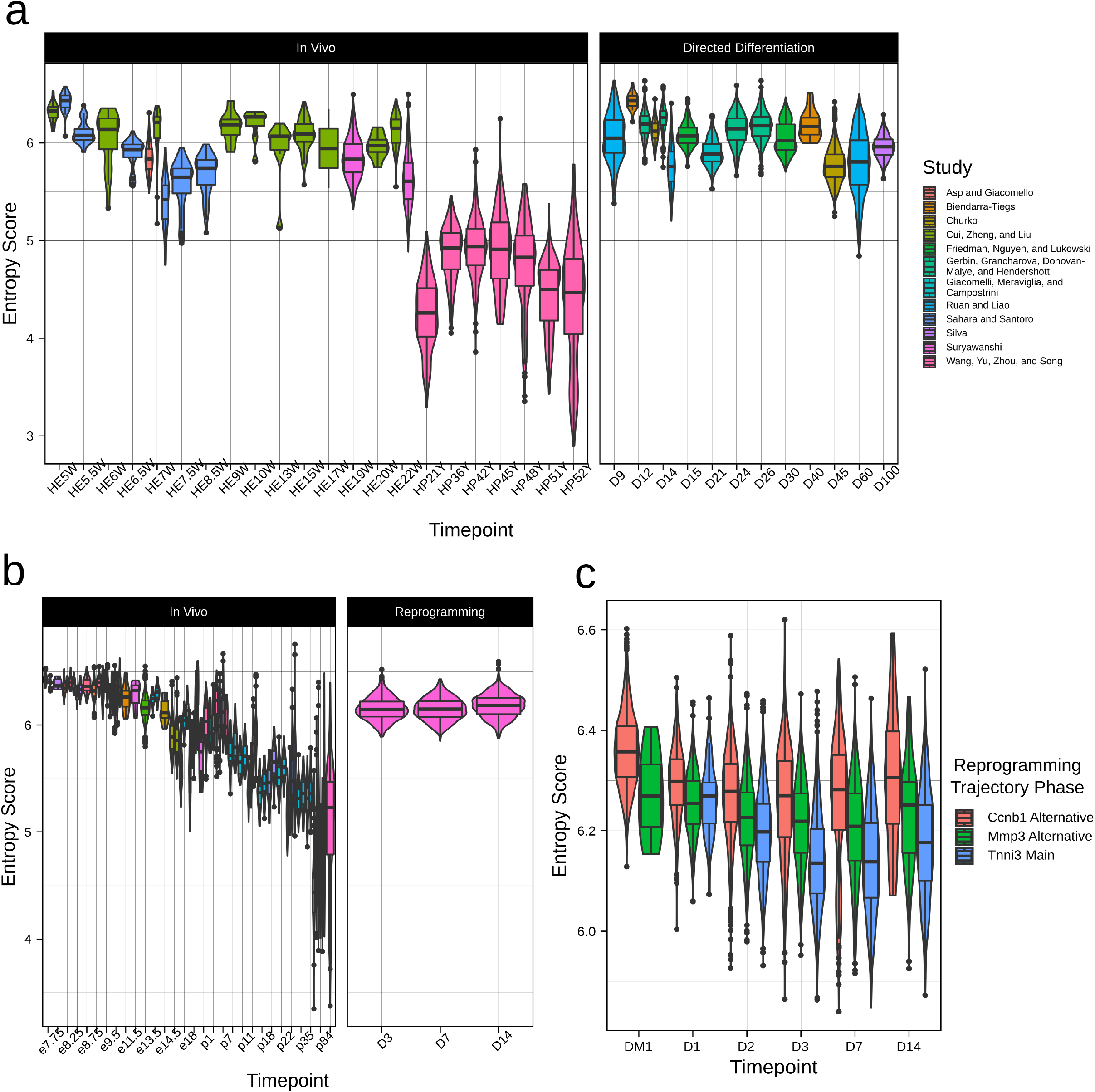
Entropy score quantifies maturation status of PSC-CMs and iCMs. **a**. Comparison in entropy score between human *in vivo* CMs and human PSC-CMs. Left side of figure reproduced from **Figure 4b. b**. Comparison in entropy between mouse *in vivo* CMs and mouse iCMs. Left side of figure reproduced from **Figure 4b. c**. Entropy score for three reprogramming pathways - a canonical Tnnt2^+^ iCM pathway and two alternative pathways (Ccnb1^+^ and Mmp3^+^).

### Reprogrammed CMs present with embryonic-like maturation status

In addition to directed differentiation of PSCs, another approach that has been explored to generate CMs *ex vivo* is direct reprogramming of fibroblasts to CM-like cells (iCMs) by various protocols (48). We used entropy score to analyze the reprogramming of mouse neonatal fibroblasts to iCMs by overexpression of Gata4, Mef2c, and Tbx5. Focusing only on cells with CM-like signature, we found that entropy score showed limited change between D3 and D14 of reprogramming (**Figure 5b**). iCMs remained at a mid-embryonic stage of maturation, comparable to e13.5-e14.5 in mouse *in vivo* CMs. Moreover, compared to PSC-CMs at the same timepoint of differentiation, iCMs displayed higher entropy. This result matches earlier findings that direct reprogramming less effectively recapitulates native gene regulatory networks compared to directed differentiation (49).

We further explored change in entropy score across multiple reprogramming pathways. The authors of the dataset identified a branching reprogramming trajectory (50). Reprogrammed cells entered either a canonical iCM route (e.g. Tnni3^+^) or two alternative pathways - one characterised by activation of Mmp3 and another marked by cell cycle progression (e.g. Ccnb1^+^). Using the authors’ annotations, we classified all cells in the dataset (including those without a CM signature) into one of these three pathways and assessed the entropy score for cells in each pathway (**Figure 5c**). At D1 of reprogramming, cells in all three pathways show similar entropy score. However, from D1 to D3, cells in the canonical iCM pathway show more notable decrease in entropy score, and indeed remain at a lower entropy score than cells in other pathways. Thus, while iCMs still present with a notably immature status compared to *in vivo*, they display some improvement in maturation status compared to cells arrested in alternative reprogramming pathways.

### Entropy score decreases over pancreatic beta cell and hepatocyte maturation

We primarily focused our attention on quantifying PSC-CM maturation, given the significant clinical need for generating CMs *ex vivo*. However, incomplete maturation and difficulty in assessing maturation status affect other tissue contexts as well. As proof of concept, we computed entropy score for *in vivo* mouse datasets of pancreatic beta cells (**Figure 6a**) and hepatocytes (**Figure 6b**). As with CMs, entropy score decreased over time for both celltypes, though with celltype-specific dynamics. For example, beta cells show a large postnatal drop in entropy score, likely corresponding to birth-related metabolic changes and need for insulin. By contrast, hepatocytes show a more steady decline in entropy score, though further datasets will be necessary to more thoroughly characterize these dynamics. Moreover, it must be noted that unlike CMs, beta cells and hepatocytes may continue to proliferate postnatally (51, 52), which may affect the interpretation of the entropy score in older tissues. Nevertheless, these preliminary results support the applicability of entropy score to non-cardiac tissue engineering contexts as well.

**Fig. 6.**
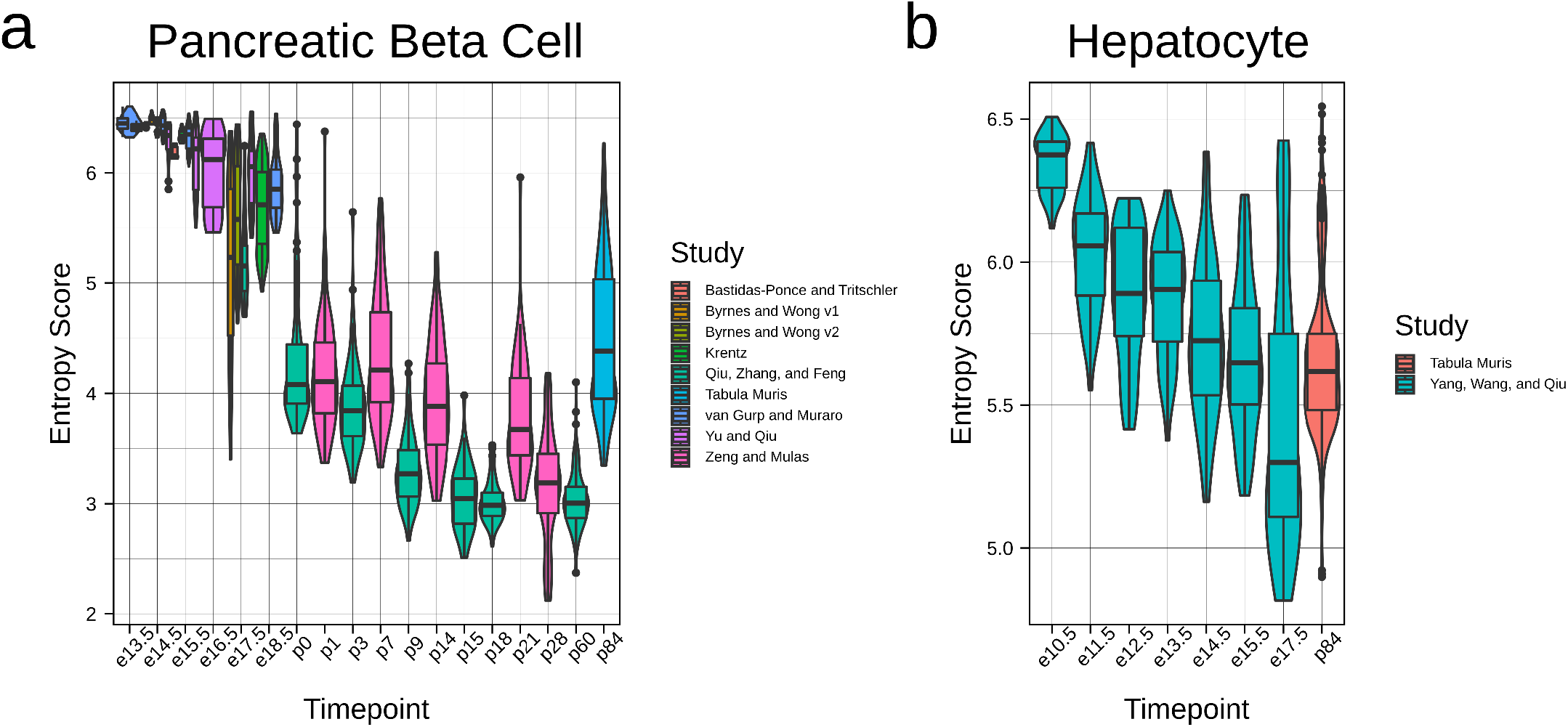
Entropy score decreases over maturation in non-CM tissue contexts. **a**. Entropy score for mouse *in vivo* pancreatic beta cells taking from publicly available datasets. **b**. Entropy scores for mouse *in vivo* hepatocytes.

## Discussion

Here, we present the use of transcriptomic entropy score for quantifying cellular maturation at the single cell level. Our approach builds on the well-known Shannon entropy to generate a metric of CM maturation from scRNA-seq data that is robust to a range of sequencing protocols and potential batch effects. In particular, entropy score enables direct benchmarking of *in vitro* PSC-CM maturation against their *in vivo* counterparts. This is particularly important because endogenous development is the gold standard for instructing PSC-derived tissue maturation. Correspondingly, we believe that perturbations to engineer maturation must be compared against this gold standard rather than an *in vitro* control. Our newly developed entropy score enables comparison of PSC-CMs against the full trajectory of endogenous CM maturation. Entropy score can thus be used to better assess PSC-CM maturation methodologies, and guide development of tissues that better recapitulate the adult CM phenotype. Moreover, while we focused on CM maturation here, we demonstrated the extensibility to other tissues as well. Given the increasing availability of both endogenous and PSC-derived scRNA-seq datasets, we expect that broad application of entropy score will enable development of improved tissues for clinical use.

It should be noted, however, that we do not see entropy score as the end-all for maturation quantification. In addition to potential discrepancies between transcript and protein level expression (53), the mature cellular phenotype encompasses numerous functional parameters that may be only partially captured at the transcriptomic level (54). We envision entropy score as complementing existing celltype-specific functional assays to advance a more complete assessment of single cell maturation status.

Through meta-analysis of over 45 scRNA-seq datasets of CMs, we were able to gain insights into the dynamics of CM maturation. In particular, we were interested to note the existence of a perinatal phase of maturation *in vivo*, initiating at ∼e16.5-e18.5, during which CM entropy score rapidly decreased. Entropy score continued to decrease until ∼3-4 weeks postnatally. We previously hypothesized the existence of a critical perinatal window for CM maturation, and postulated that disruption of this window *in vitro* leads to maturation arrest (46). The significant decrease in entropy observed in our study supports the perinatal window hypothesis. Moreover, late-stage PSC-CMs remained arrested at an entropy score similar to those of e16.5 CMs *in vivo*. To date, mechanistic understanding of PSC-CM maturation arrest has been limited, but may involve progressive disruption of cardiac gene regulatory networks (18). Our results suggest that focus should be placed on trying to understand regulators of perinatal maturation *in vivo*, and determining discrepancies in activity of these regulators *in vitro*. Whether similar mechanisms underlie maturation arrest in other PSC-derived cells remains a question for further studies.

In this study, we found that entropy score could be applied to scRNA-seq datasets generated from a wide range of protocols. Excluding the quality control steps, entropy score is computed from information in one cell at a time, independent of other cells or datasets. Nevertheless, entropy score shows strong concordance with CM maturation status in a comparable manner across dataset. This is particular novel as, thus far, direct comparisons across studies has been limited by confounding batch effects. Moreover, current batch correction algorithms may be poorly suited to integration of datasets along a continuous trajectory. Additionally, scaling batch correction algorithms to many datasets may be complex and computationally intensive. By contrast, entropy score has limited computational demands and can scale easily to allow for comparison of many datasets.

We were particularly intrigued to note the comparability of entropy scores across datasets with entirely different datatypes (e.g. reads, UMIs). For example, it is well known that PCR amplification in scRNA-seq protocols can lead to biases (55), which was one of the motivations for the development of UMIs. However, entropy scores were comparable for UMI datasets prior to and after collapsing UMIs. Likewise, datasets generated from full-length protocols did not display notable biases in entropy score. This observation may have been incidental to the datasets we studied - for example, high quality datasets may have presented with sufficiently low amplification bias to enable comparison. It is possible that entropy score is less robust to more extreme cases of amplification bias. We do not believe our finding precludes the use of best practices for scRNA-seq protocols, including the use of UMIs for many experimental designs. Nevertheless, we were encouraged that entropy score could be used to facilitate cross-comparison between otherwise incompatible datatypes.

One technical limitation of entropy score was its poor performance with Drop-seq datasets. We consistently found that Drop-seq datasets presented with higher entropy than data generated at similar timepoints through other protocols. This may be a consequence of depth; the Drop-seq datasets that we tested were the lowest depth studies tested and below our identified optimal depth threshold. However, given the increasing prevalence of other high-quality droplet-based protocols (in particular, 10x Chromium), we believe this is not a major limiting factor to the use of entropy score. We additionally did not test single nuclear RNA-seq datasets, both due to concerns of depth and because we expected that the gene distribution would be inherently different from whole cell studies (56). Nevertheless, the emergence of methods for isolation of whole adult CMs in mouse and human (31, 57, 58) may reduce the future need for nuclear RNA-seq.

At the single cell level, CM maturation proceeds heterogeneously along a unidirectional trajectory (33). We were therefore curious to know the extent to which entropy score could capture single cell positioning along this trajectory, in effect functioning as a pseudotime metric. Entropy score only modestly correlated with other established pseudotime methods, though all methods recovered similar differentially expressed genes. These discrepancies may be due to transcriptomic noise in single cell data. However, it should be emphasized that entropy score works in a fundamentally different manner than many trajectory inference methods. Most trajectory inference methods utilize some type of dimensionality reduction step prior to curve fitting. By contrast, outside of the subselection of highly expressed genes, entropy score uses no dimensionality reduction step. Moreover, entropy score makes no assumptions about relationships between cells - all relevant information is calculated independently for each cell. Despite being agnostic to cell-cell relationships, entropy score accurately captures CM maturation expression trends. Commonly used dimensionality reduction methods have been shown to distort local neighbourhoods and affect trajectory reconstruction (59), and thus entropy score may more optimally capture single CM dynamics in maturation.

Entropy score has several important antecedents that must be acknowledged. Our work is similar to StemID (25), which uses Shannon entropy to assign progenitor state within a trajectory. We extend this usage with several gene filtering steps to better facilitate cross-study comparison. Shannon entropy is also utilized in SLICE (27), which computes entropy based on functional annotations of genes, and SCENT (29), which computes entropy within a protein-protein interaction network. Both approaches are powerful for constructing trajectories for differentiating cells. However, unlike differentiation, CM maturation is characterized by continuous rather than step-wise or switch-like changes. For this purpose, an entropy score built directly on gene expression levels is both simpler to compute and more appropriate. Lastly, our work is similar conceptually to CytoTRACE (26), which leverages gene diversity to order cells by differentiation status. Directly comparing number of genes expressed by each cell is confounded by cross-study differences in depth and sensitivity, however. CytoTRACE addresses this by using a smoothing step within dataset. However, this limits its use for datasets with few cells or representing fewer maturation states. By contrast, outside of quality filtering, entropy score performs computations on each cell independently, extending its utility to more datasets. We believe these differences improve the utility of entropy score for benchmarking the maturation status of PSC-derived tissues.

### Materials and Methods

All methods, including wet lab and computational methods, can be found in the Supplementary Information. Raw data for the maturation reference can be found on GEO at GSE147807. Code to generate figures in this manuscript as well as the counts tables for the datasets analyzed in this manuscript can be found on Github at https://github.com/skannan4/cm-entropy-score.

## ACKNOWLEDGMENTS

We thank Dr. Deborah Andrew for allowing us to use her group’s COPAS LP-FACS instrument. Sequencing experiments were done through the Johns Hopkins Transcriptomics and Deep Sequencing Core, with the help of Dr. Haiping Hao, Linda Orzolek, Dr. Jasmeet Sethi, and Kelly Laughlin. We additionally thank Dr. Marc Halushka, Dr. Patrick Cahan, Stephanie Yang, and Yuqi Tan for helpful comments in preparing this manuscript. This work was supported by grants from NICHD/NIH, AHA, and MSCRF.

The authors have no competing interests to declare.

## Supplementary Information for

### Supporting Information Text

Shannon entropy has had long-standing applications in developmental biology as well as transcriptional analysis (1**?**). A standard form for Shannon entropy *S* is:

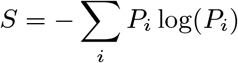

where *P*_*i*_ represents individual probabilities for events of interest. Here, we define *P*_*i*_ as the probability of selecting a given gene *i* in a cell. From scRNA-seq data, this can be computed by simply dividing the number of counts for gene *i* by all of the gene counts in a given cell. For our entropy score, we similarly use Shannon entropy, except after subsetting the top 1000 highest expressed genes to enable sensitivity control.

### Supplementary Methods

#### Mice

To generate mice for our reference dataset, we crossed B6.FVB-Tg(Myh6-cre)2182Mds/J mice (aMHC-cre, Jackson Laboratory, Stock No. 011038) with B6.129(Cg)-Gt(ROSA)26Sor^tm4(ACTB-tdTomato,-EGFP)Luo/J^ (mTmG, Jackson Laboratory, Stock No. 007676). Both mice have C57BL/6J congenic background. All animals were maintained compliant to protocols by the Johns Hopkins Animal Care and Use Committee.

#### CM Isolation

For isolation of CMs from e14-p4 timepoints, we used the neonatal cardiomyocyte isolation kit from Miltenyi Biotec in conjunction with the gentleMACS Dissociator. For later timepoints, we performed Langendorff isolation of CMs. We prepared the following buffers:

- Perfusion buffer: 120 mM NaCl, 5.4 mM KCl, 1.2 mM NaH_2_PO_4_, 20 mM NaHCO_3_, 5.5 mM glucose, 5 mM BDM, 5 mM Taurine, and 1 mM MgCl_2_, adjusted to pH 7.4
- Digestion buffer: 40 mL Perfusion buffer plus 35.8 mg Collagenase Type II (Worthington CLS-2), 3 mg Protease (Sigma P5147)
- Tyrode’s buffer: 140 mM NaCl, 5 mM KCl, 10 mM HEPES, 5.5 mM glucose, and 1 mM MgCl_2_, adjusted to pH 7.4

We used a horizontal (i.e. non-hanging) Langendorff apparatus with a chamber filled with perfusion buffer. To perform isolation, we first performed isofluorane anaesthesia on non-heparinized mice. Mice were observed until clearly anaesthetized and unresponsive to toe pinch, and subsequently euthanized by cervical dislocation. The heart was then rapidly excised from the chest and cannulated to the Langendorff apparatus. Flow time and rate of flow were dependent on the age of the mouse and were typically judged based on completeness of digestion to touch. Subsequently, the left ventricular free wall was excised and minced. We filtered isolated cells through a 100 *µ*M screen to eliminate large tissue chunks, spun down at 800 RPM for 1 minute (Eppendorf centrifuge 5702), and resuspended cells in 10 mL Tyrode’s buffer.

#### LP-FACS

We have detailed our LP-FACS approach previously (2). We reproduce our methods here. We utilized a COPAS SELECT instrument (Union Biometrica). The COPAS SELECT was updated and rebranded as the FP-500, but the protocol here study does not use the new features and thus the two are functionally indistinguishable. We optimized sorting for cardiomyocytes by using a sort delay of 8 and sort width of 6. Additionally, we used the following fluorescence settings: ext gain 50, green gain 200, yellow gain 200, red gain 255, extension integral gain 50, green integral gain 200, yellow integral gain 200, red integral gain 255, green PMT 800, yellow PMT 800, red PMT 1100. Coincidence check was selected to ensure proper single event sorting. We typically flowed cells between 20 - 60 events/second. We maintained cells in Tyrode’s buffer during the sort and sorted them into Tyrode’s buffer. To run the machine, we used ClearSort Sheath Fluid (Sony, Lot 1218L345).

#### scRNA-seq Library Preparation and Sequencing

We performed sequencing using the mcSCRB-seq protocol (**?**). The protocol has been described at protocols.io at dx.doi.org/10.17504/protocols.io.p9kdr4w. Pooled libraries were sequenced on one mid-output lanes of the Illumina NextSeq500 with 16 base pair barcode read, 8 base pair i7 index read, and 66 base pair cDNA read design.

#### Computational Analyses

All analyses performed in the paper were done in R; code to reproduce the figures can be found at our Github (https://github.com/skannan4/cm-entropy-score). Dataset characteristics are presented in **Supplementary Tables 1** and **2**, and details of each individual dataset are described in the **Appendix**. Dimensionality reduction as well as dataset integration for **Figure 1** was done using Monocle 3. Differential gene expression analysis for **Figure 4** was done using Monocle 2, replacing Monocle 2’s generated pseudotime with entropy score or pseudotime from other methods as appropriate.

### Supplementary Figures

Supplementary figures for the manuscript can be found below.

**Fig. S1.**
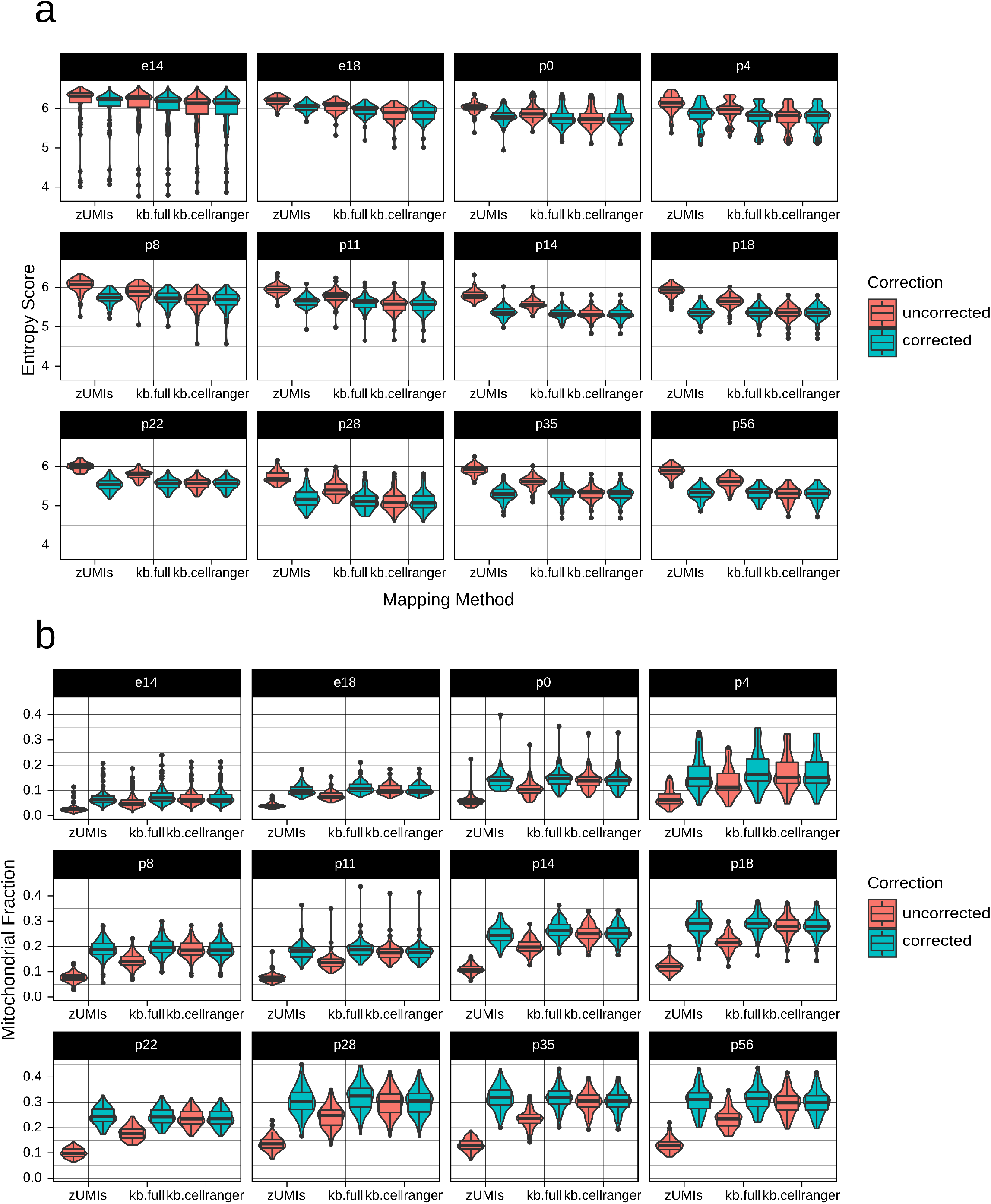
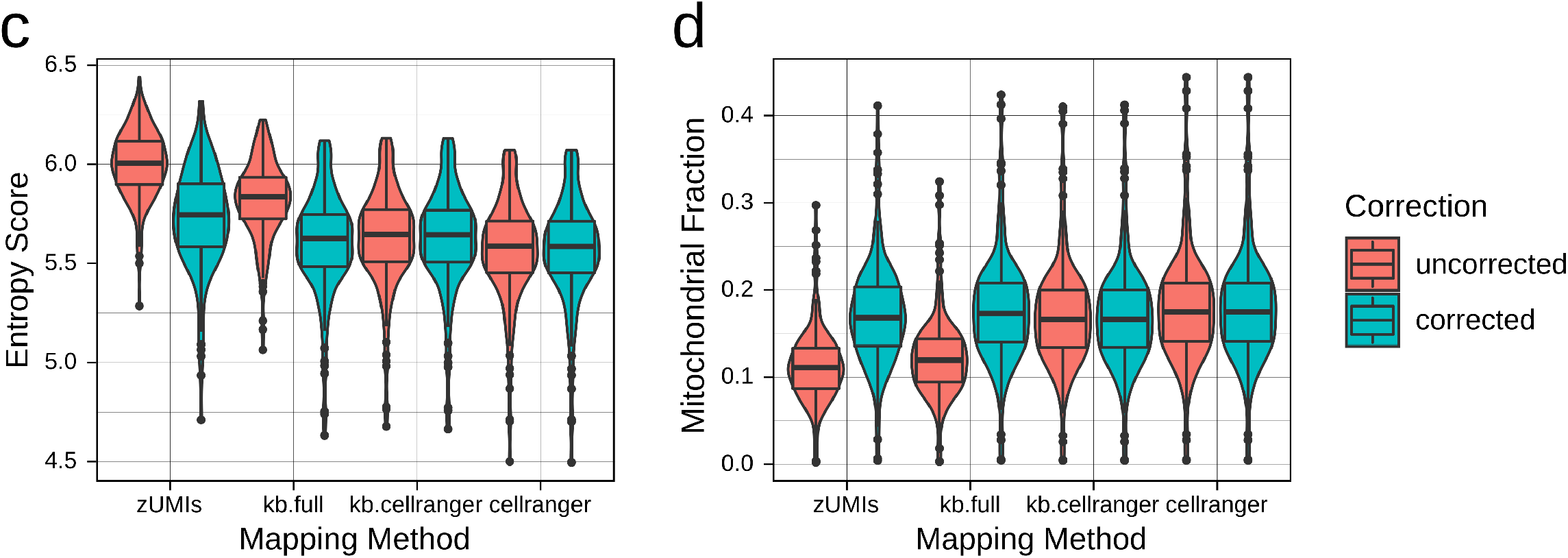
Correction of mitochondrial pseudogenes enables consistent entropy score measurements across mapping/counting pipelines. **a**. Entropy scores for the maturation reference dataset mapped by zUMIs, kallisto|bustools with the full reference, and kallisto|bustools with the CellRanger reference. Pre-s and post-correction scores are shown. **b**. As in **a**, showing mitochondrial proportions. **c.** Entropy scores for the 10x Chromium heart dataset mapped by zUMIs, kallisto|bustools with the full reference, kallisto|bustools with the CellRanger reference, and CellRanger. **d.** As in **c**, showing mitochondrial proportions.

**Fig. S2.**
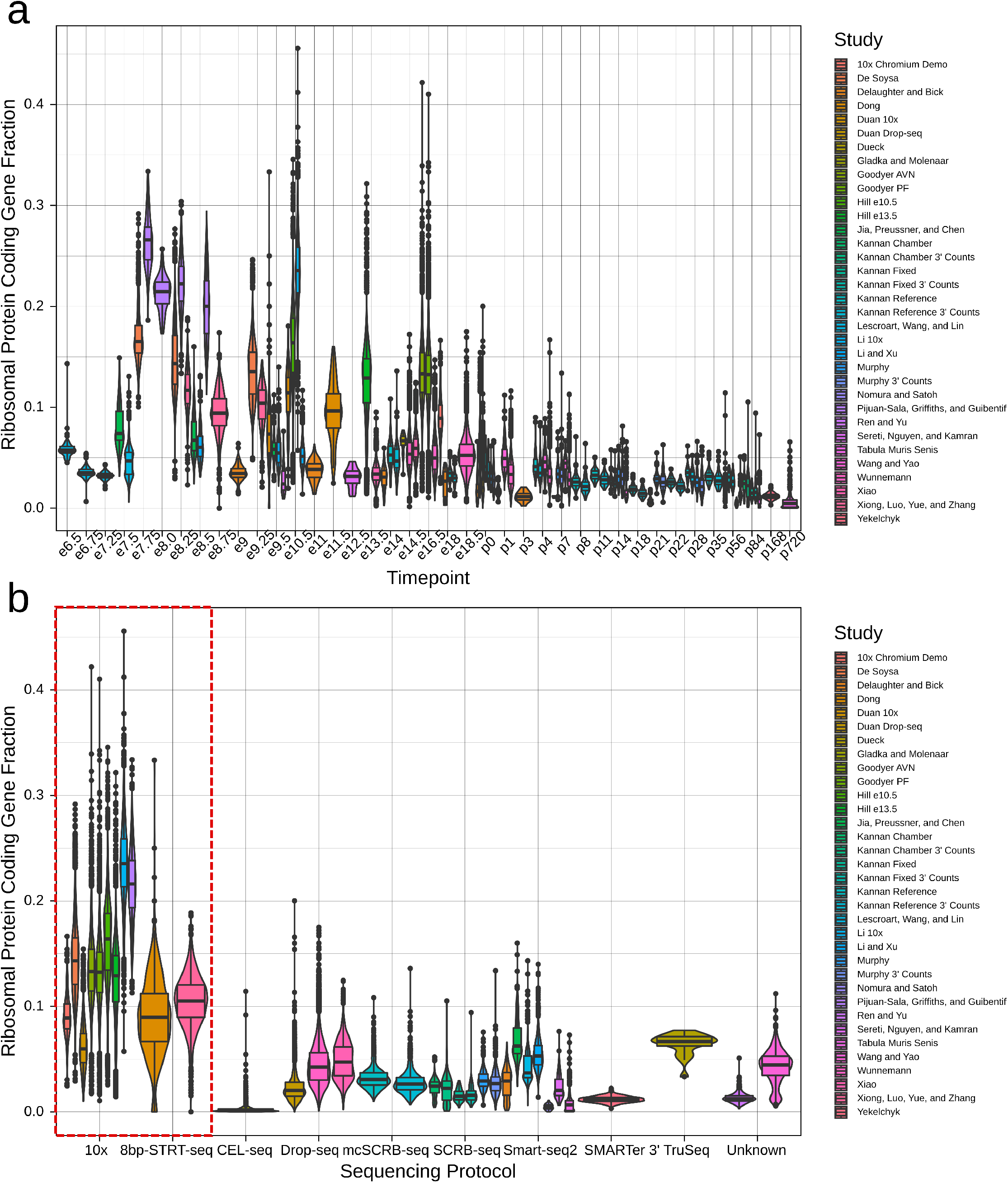
Ribosomal protein-coding genes are expressed in a sequencing protocol-specific manner. **a**. Proportion of ribosomal protein coding genes in mouse *in vivo* datasets, grouped by timepoint. **b**. Proportion of ribosomal protein coding genes in mouse *in vivo* datasets, grouped by library preparation method. 10x v1-v3 protocols have been coalesced together for the purposes of this figure.

**Fig. S3.**
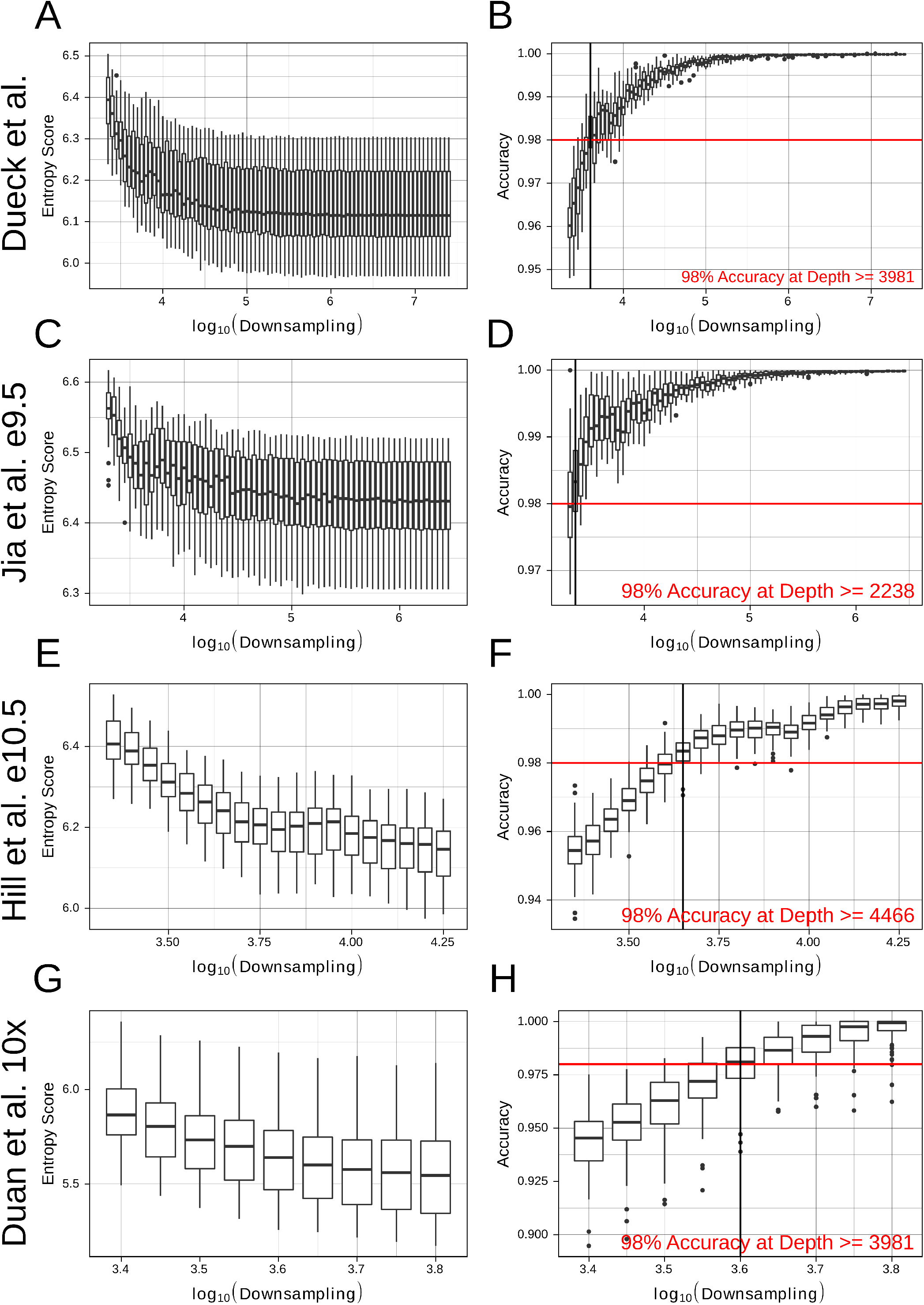
Entropy score is robust across a range of sequencing depths. For each of four datasets, we performed subsampling and computed the entropy score as well as accuracy (calculated as deviation from baseline entropy score). At each stage, we included only cells with genes > 1000, and subsampled only to a depth where the median number of genes remained > 1000. Data is shown for **a-b**. Dueck et al. **c-d**. Jia et al. at e9.5. **e-f**. First 100 cells from Hill et al. at e10.5. **g-h**. First 100 cells from Duan et al.

**Fig. S4.**
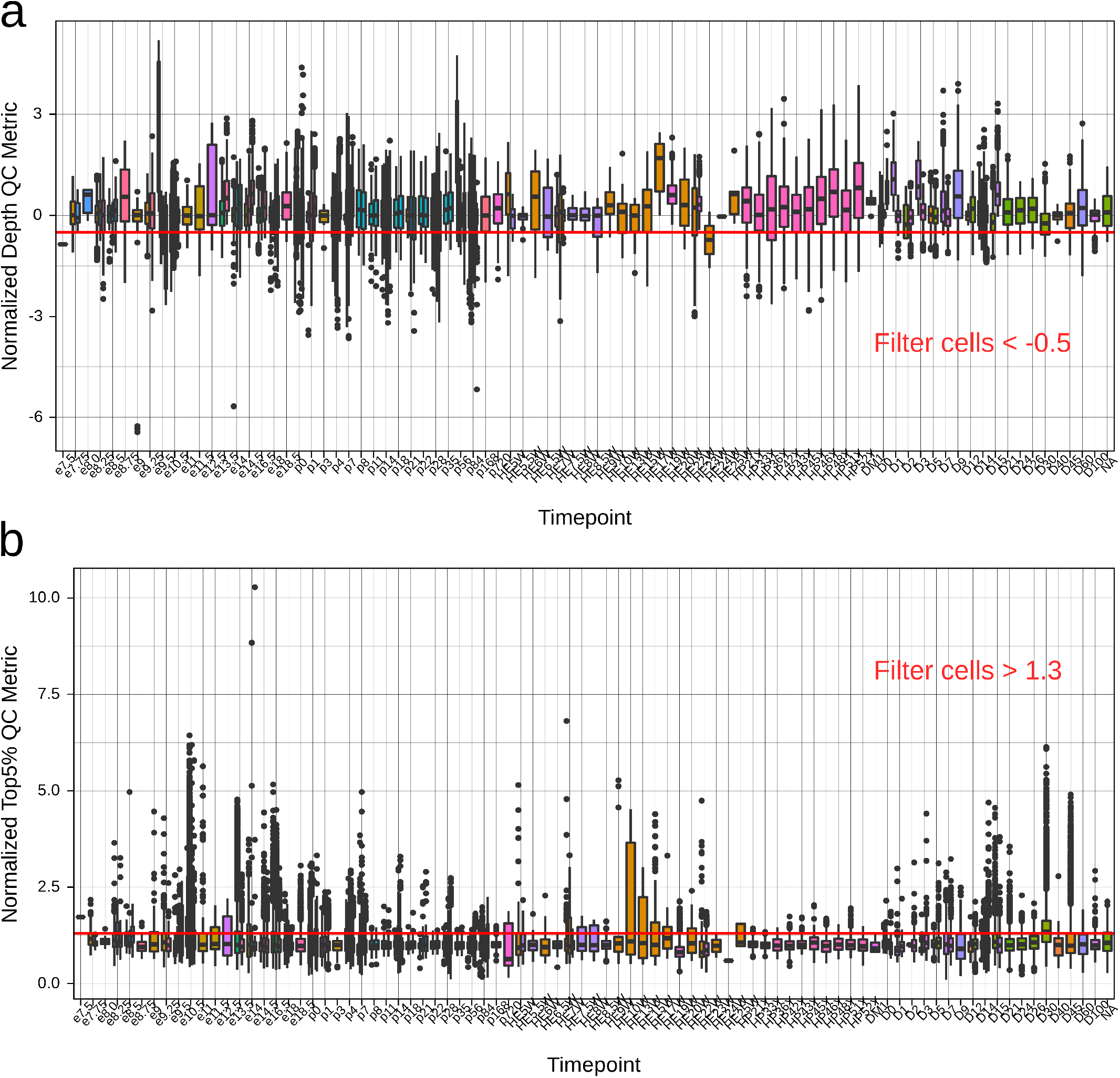
Poor quality single cells can be identified and removed with normalized depth and top 5 gene percentage metrics. **a**. Normalized depth QC metric for all datasets. Red line indicates the threshold of −0.5. **b**. Normalized top 5 gene percentage metric for all datasets. Red line indicates the threshold of 1.3.

**Fig. S5.**
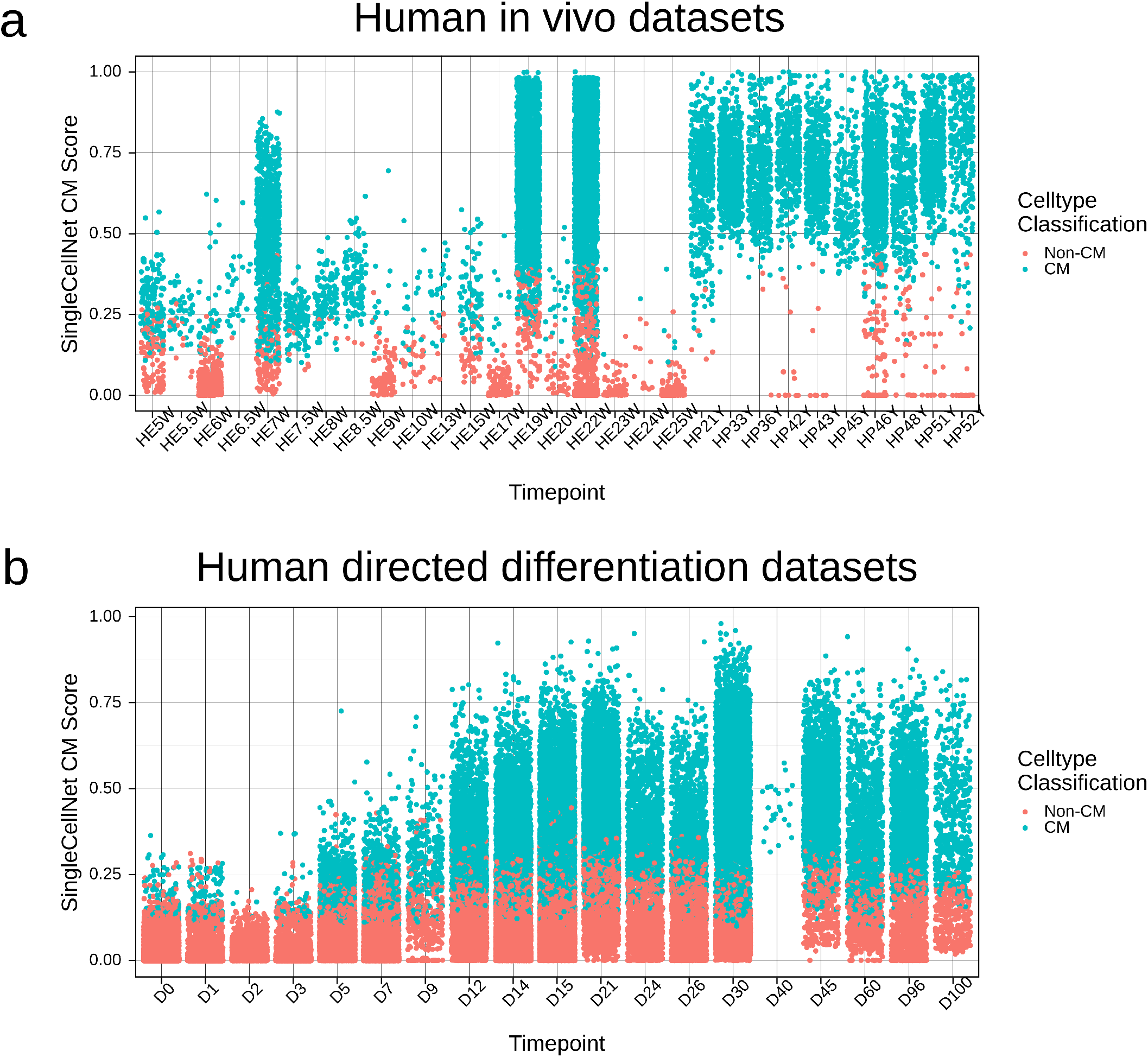
SingleCellNet identifies single cells with CM signature. Cells are labeled based on whether their highest classification was for “cardiac muscle” or another celltype. **a**. For human *in vivo* datasets. b. For human *in vitro* directed differentiation datasets.

**Fig. S6.**
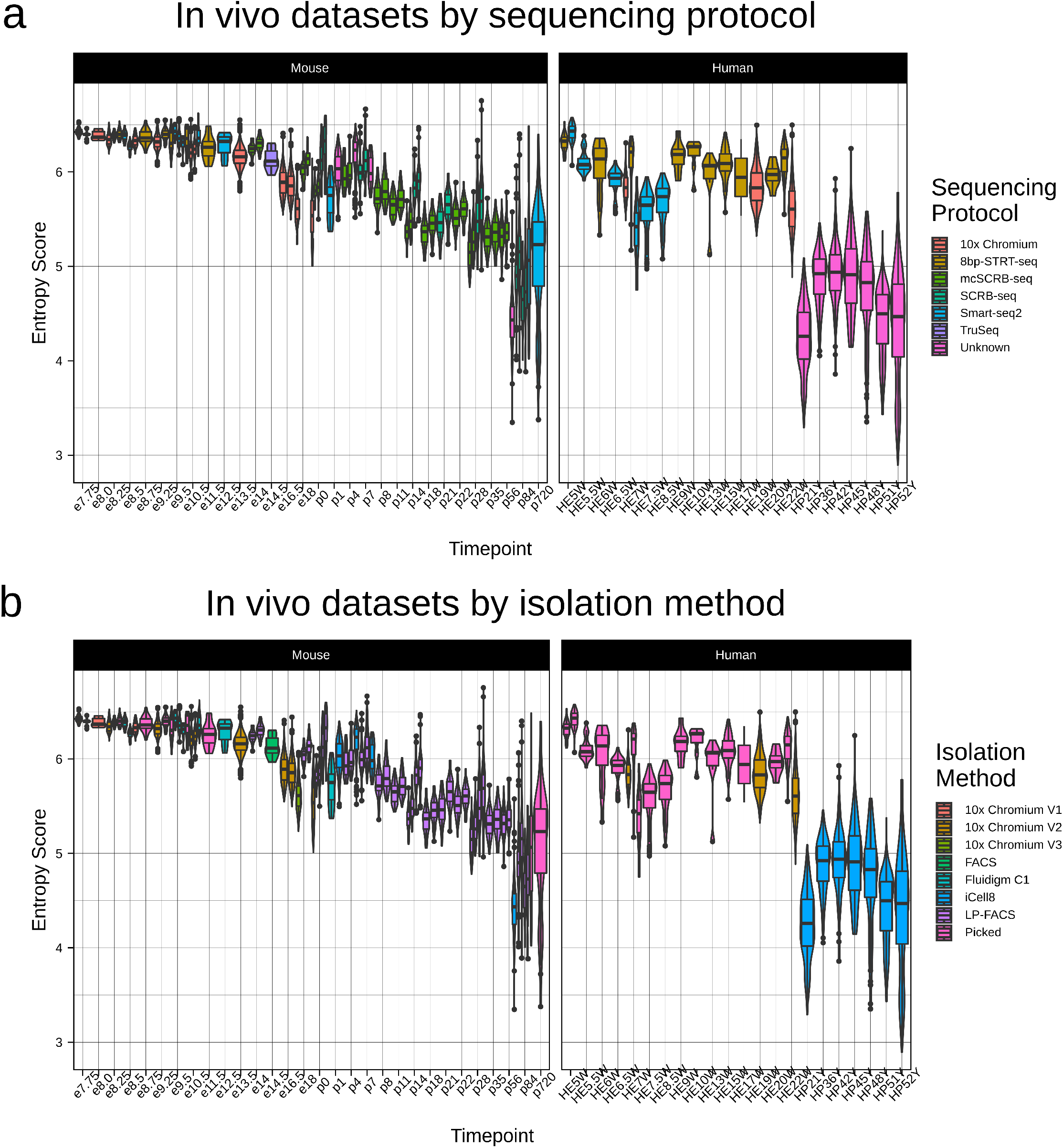
Entropy score enables comparison of maturation status of CMs from scRNA-seq datasets with diverse characteristics. This figure corresponds to **Figure 2b**, but with boxplots coloured by **a**. sequencing protocol and **b**. isolation method.

**Fig. S7.**
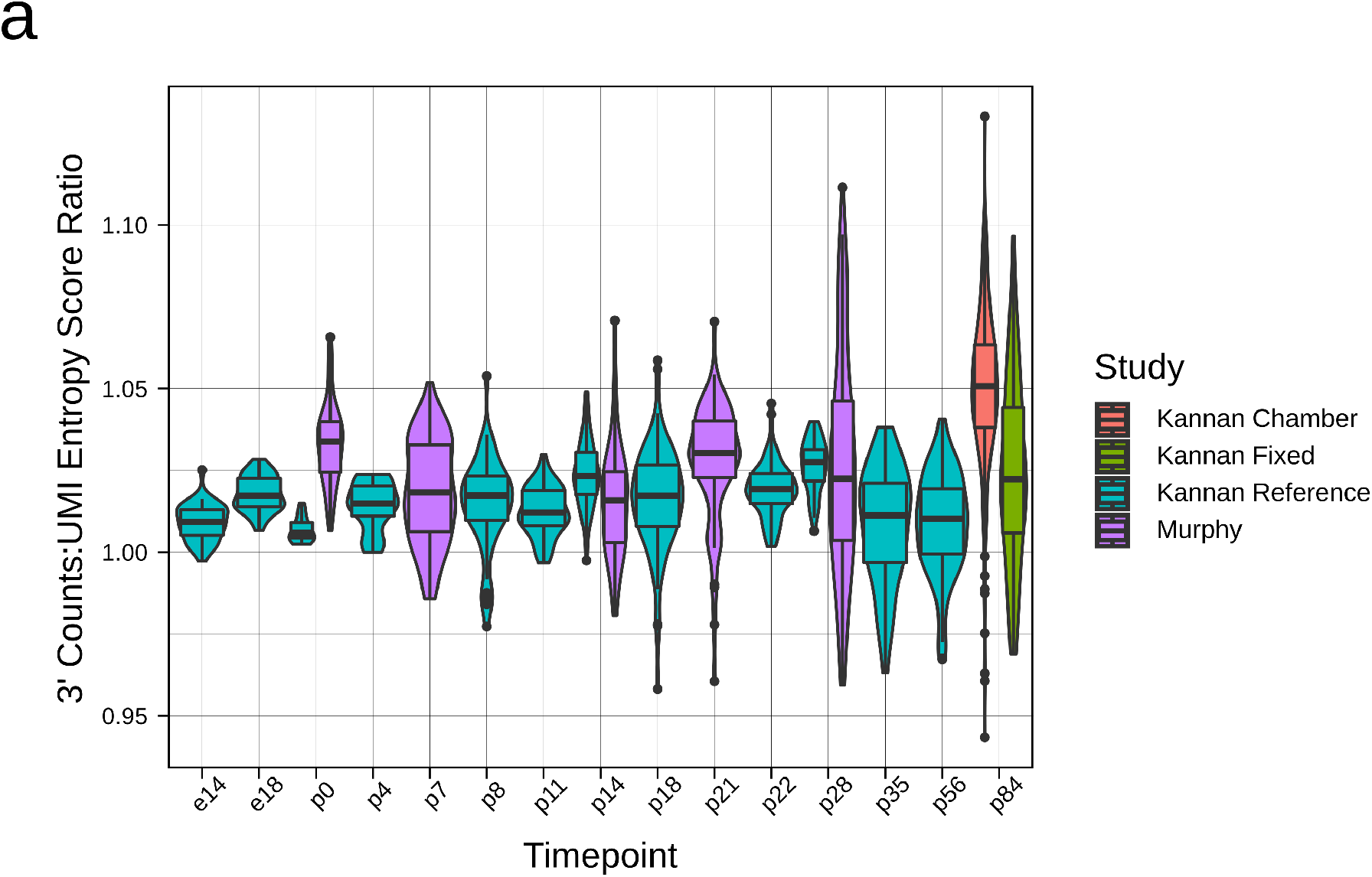
Entropy score is consistent for UMI datasets pre- and post-UMI collapsing. **a**. Ratio of entropy score for UMI datasets computed prior to vs. after UMI collapsing.

**Fig. S8.**
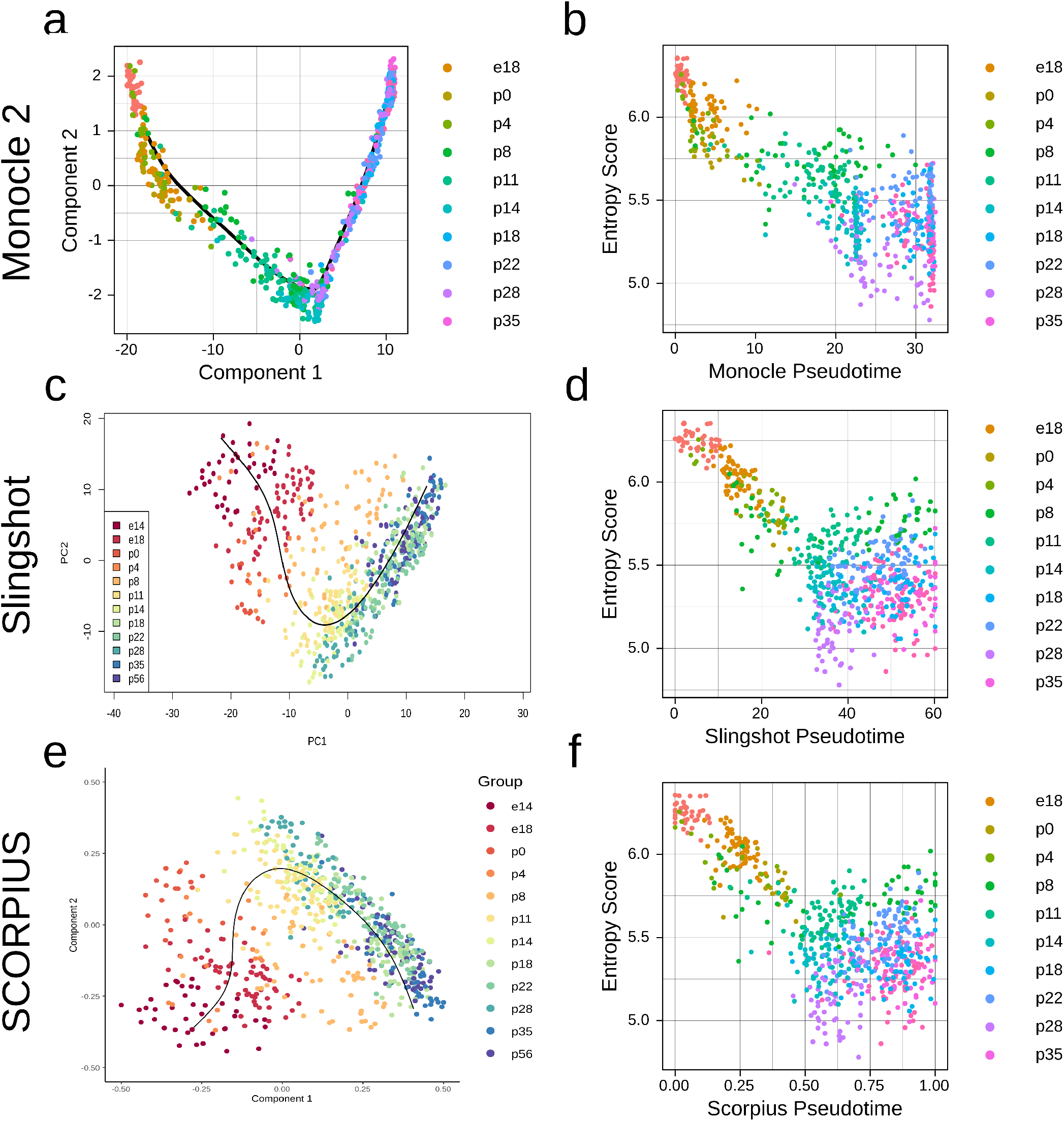
Entropy score correlates modestly with previous trajectory inference methods. We reconstructed trajectories of our maturation reference dataset using **a-b**. Monocle 2, **c-d**. Slingshot, and **e-f**. SCORPIUS.

**Fig. S9.**
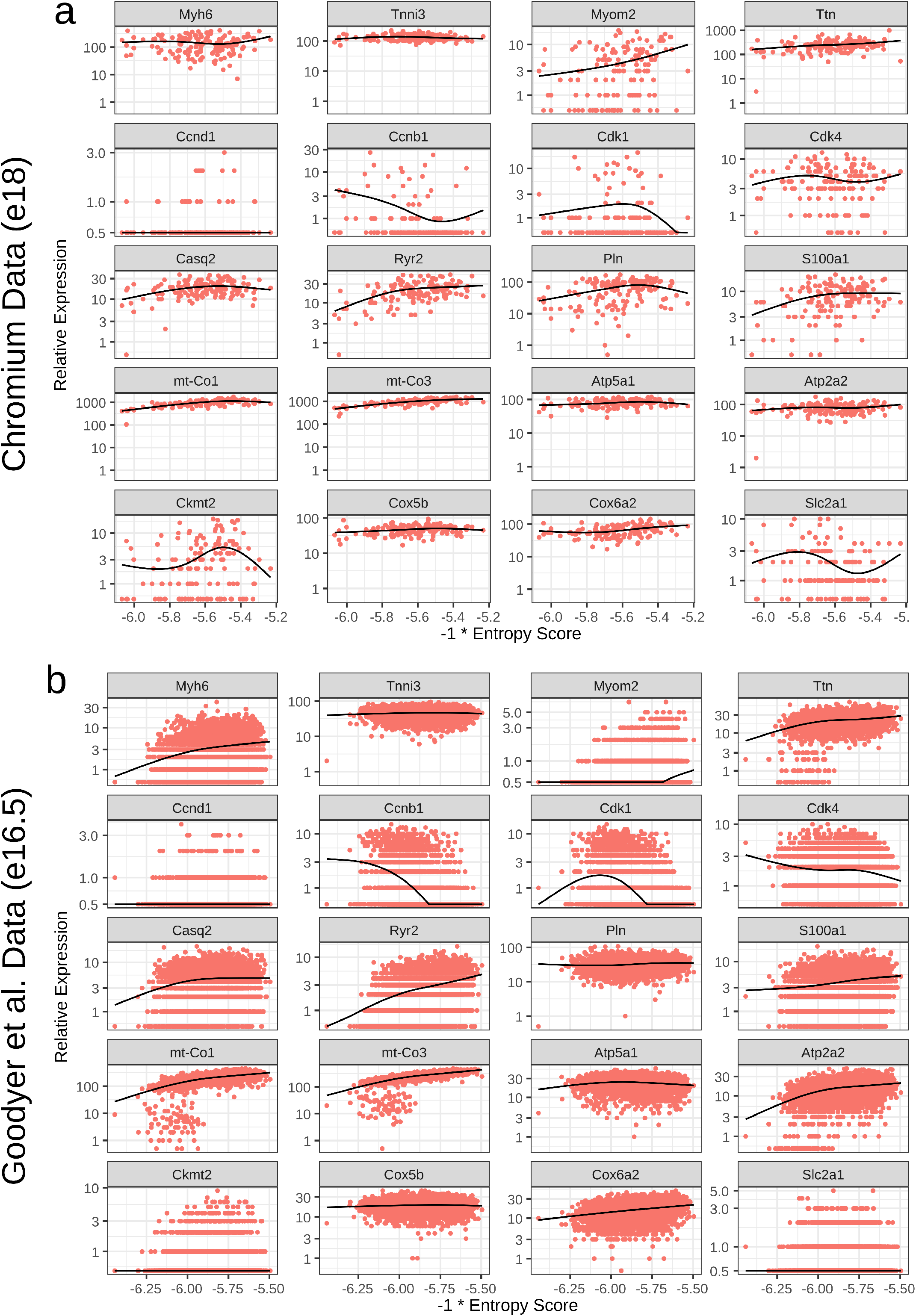
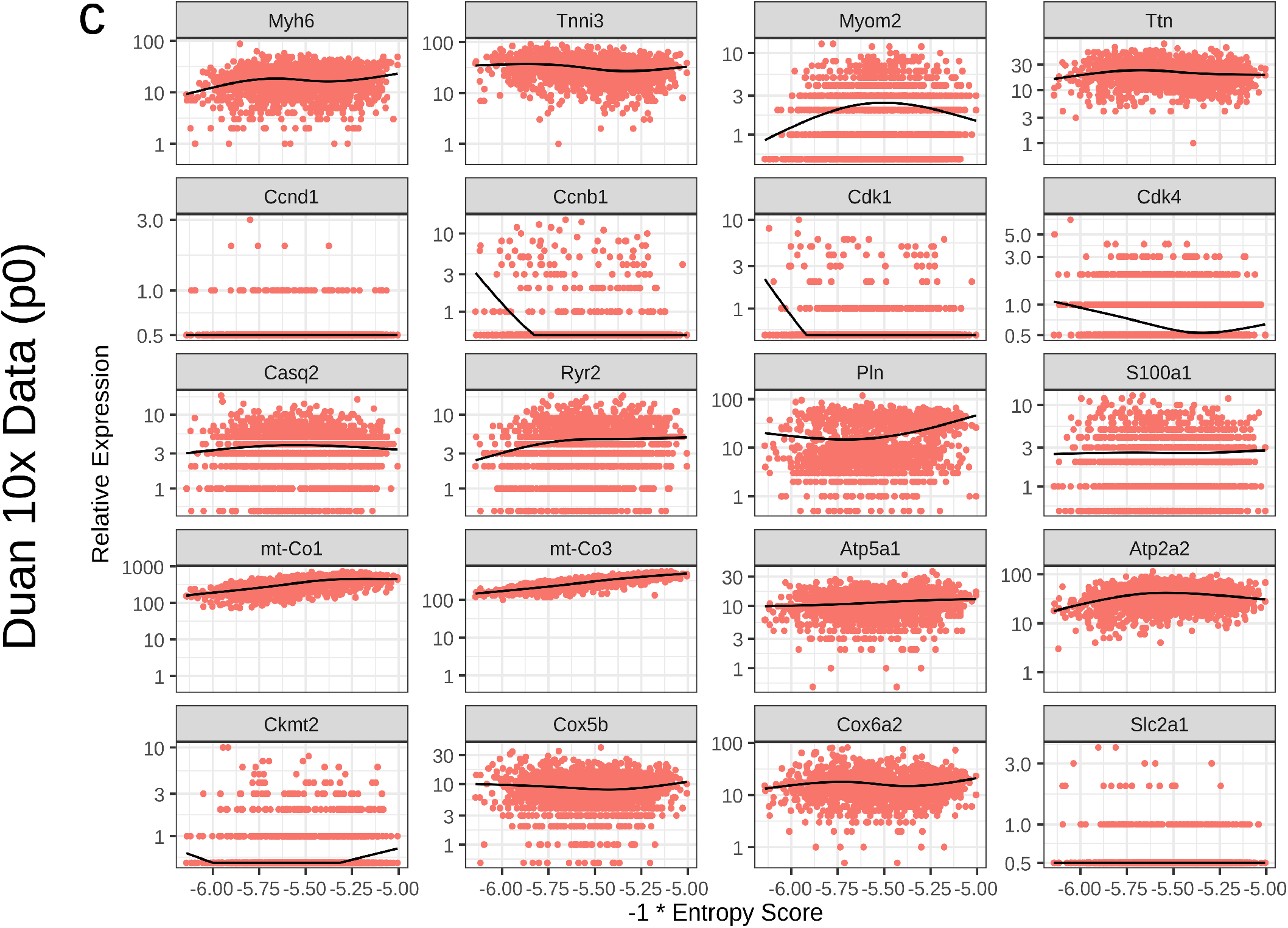
Entropy score captures CM maturation-related gene expression trends in one-timepoint datasets. Gene trends across entropy score, as in **Figure 3c**, are plotted for **a**. 10x Chromium heart dataset, **b**. Goodyer et al., and **c**. Duan et al. Gene trends across entropy score, as in **Figure 3c**, are plotted for c. Duan et al. Suraj

## Appendix

This appendix contains information about all of the datasets included for analysis in the study, including how they were acquired and processed. Metrics for each dataset are provided in **Supplementary Tables 1** and **2**. We have relevant data on Github at https://github.com/skannan4/cm-entropy-score. This includes an R workspace containing counts table for every dataset in this study, an R workspace containing just the final processed calculations (for lower memory usage), code for functions relevant to usage of entropy score, and code to reproduce the figures in the manuscript. If any additional information is required, we encourage direct inquiries and aim to respond as soon as possible.

### Mouse In Vivo

*10x Chromium*

*10k Heart Cells from an E18 mouse (v3 chemistry)*

We downloaded the filtered feature/cell matrix from the 10x chromium datasets website (https://support.10xgenomics.com/single-cell-gene-expression/datasets/3.0.0/heart_10k_v3?). We subsequently performed UMAP + clustering using Seurat and selected the healthy cardiomyocytes as the cluster clearly expressing cardiomyocyte markers and having comparable/higher read/gene counts to other clusters.

*T. Yvanka de Soysa et al. (Casey Gifford, Deepak Srivastava)*

*Single-cell analysis of cardiogenesis reveals basis for organ-level developmental defects*

We pulled the UMI data from the tables uploaded by the authors at GEO (GSE126128), selecting only wild-type cells at all available timepoints. We used the annotations provided by the authors as the source data to Extended Data Figure 1 to identify tissue, and selected out cells labeled as “myocardium.” We used the annotations provided by the authors as source data to Extended Figure 1 to annotate myocardial region, though we did not filter based on these annotations.

*Daniel DeLaughter and Alexander Bick et al. (Jonathan Seidman, Christine Seidman)*

*Single-Cell Resolution of Temporal Gene Expression during Heart Development*

We pulled the FastQ data from the authors’ private database (https://b2b.hci.utah.edu/gnomex/) and mapped using STAR/FeatureCounts. We then performed T-SNE + clustering through Seurat and selected healthy cardiomyocytes as the clusters clearly expressing cardiomyocyte markers. However, we found that this data consistently had higher mitochondrial percentage at almost every timepoint compared to other datasets (most notably in the perinatal and postnatal timepoints). Thus, we discarded this dataset.

*Ji Dong et al. (Fuchou Tang)*

*Single-cell RNA-seq analysis unveils a prevalent epithelial/mesenchymal hybrid state during mouse organogenesis*

The uploaded data of the authors does not contain mitochondrial reads. Therefore, we pulled and mapped the FastQ data from the four heart samples from ENA (PRJNA343327) using the FASTerQ approach with Kallisto|Bustools. Read 1 was used as the cdna read (base pairs 45 to 140, to avoid potential adaptor and poly A tails), while Read 2 was used for barcode (first 8 bp) and UMI (second 8 bp). Because of the short UMI length, we found that Kallisto|Bustools discarded many UMIs during counting. We thus used a custom script to count UMIs from the Kallisto|Bustools output.

*Jialei Duan et al. (Nikhil Munshi, Gary Hon)*

*Rational Reprogramming of Cellular States by Combinatorial Perturbation*

The UMI data was pulled from the tables uploaded by the authors at GEO (GSE117795). We collated all of the in vivo samples, with the 10x and Drop-seq data being handled separately. We then performed UMAP + clustering through Seurat and selected healthy cardiomyocytes as the clusters clearly expressing cardiomyocyte markers while having comparable read counts to other clusters and comparable mitochondrial percentage (discarding clusters with notably high mitochondrial percentage or low genes). We could readily distinguish atrial and ventricular myocytes using myosin light chain isoforms, and annotated the cells accordingly. We found that the Drop-seq data had unusually high entropy, something we observed across multiple Drop-seq datasets (and perhaps owing to low depth); thus, we discarded the Drop-seq data.

*Hannah Dueck et al. (Junhyong Kim)*

*Deep sequencing reveals cell-type-specific patterns of single-cell transcriptome variation*

The uploaded data of the authors does not contain mitochondrial reads. Therefore, we pulled the FastQ data from the cardiomyocyte samples from ENA (PRJNA244374) and then mapped with Kallisto (pseudo in batch mode with the -quant flag).

*Monika Gladka and Bas Molenaar et al. (Eva von Rooij)*

*Single-Cell Sequencing of the Healthy and Diseased Heart Reveals Cytoskeleton-Associated Protein 4 as a New Modulator of Fibroblasts Activation*

Counts tables were kindly provided by the authors. We used UMAP + clustering through Seurat to select cardiomyocytes based on clusters expressing cardiomyocyte markers. We observed, however, that the dataset had very high mitochondrial percentages, often close to 90%. We thus discarded this dataset.

*William Goodyer (Sean Wu)*

*Transcriptomic Profiling of the Developing Cardiac Conduction System at Single-Cell Resolution*

We pulled the UMI data from tables uploaded by the authors at GEO (GSE132658). The authors kindly provided us with the clustering used in the manuscript, which we used to select out cardiomyocytes. We used the PF (left and right) and AVN datasets as the metadata was not available for the SAN data.

*Matthew Hill et al. (James Martin)*

*A cellular atlas of Pitx2-dependent cardiac development*

The UMI data was pulled from the tables uploaded by the authors (GSE131181), and the metadata tables were pulled from the same source. We used the authors’ generated clusters and selected clusters with high expression of cardiomyocyte markers. We subsequently selected only control cells from both timepoints.

*Guanshuai Jia, Jens Preussner, and Xi Chen et al. (Thomas Braun)*

*Single cell RNA-seq and ATAC-seq analysis of cardiac progenitor cell transition states and lineage settlement*

The counts data was pulled from the authors’ Github (https://github.com/loosolab/cardiac-progenitors). We used the data from both Isl and Nkx GFP lines.

*Suraj Kannan et al. (Chulan Kwon)*

*Large Particle Fluorescence-Activated Cell Sorting Enables High-Quality Single-Cell RNA Sequencing and Functional Analysis of Adult Cardiomyocytes*

This data was generated at our lab and is available at GEO (GSE133640). We used both the multi-chamber study and the lived/fixed study and included all cells from both studies. Both 3’ counts and UMIs (output from zUMIs, using intronic and exonic reads) were used for analysis.

*Suraj Kannan et al. (Chulan Kwon)*

*Transcriptomic entropy quantifies cardiomyocyte maturation at single cell level*

This data (described in this manuscript) was generated at our lab and is available at GEO (GSE147807). Both 3’ counts and UMIs (output from zUMIs, using intronic and exonic reads) were used for anaysis.

*Fabienne Lescroart, Xiaonan Wang, and Xionghui Lin et al. (Cedric Blanpain)*

*Defining the earliest step of cardiovascular lineage segregation by single-cell RNA-seq*

We download the counts data from the author’s private website (http://singlecell.stemcells.cam.ac.uk/mesp1#data). We subsequently selected only wild-type cells.

*Guang Li and Adele Xu et al. (Sean Wu)*

*Transcriptomic Profiling Maps Anatomically Patterned Subpopulations among Single Embryonic Cardiac Cells*

We pulled the counts data uploaded by the authors at GEO (GSE76118). We subsequently performed TSNE + clustering through Seurat and selected clusters clearly identifiable as cardiomyocytes by marker gene expression. We selected all wild-type cells, including from the Nkx experiment.

*Guang Li et al. (Sean Wu)*

*Single cell expression analysis reveals anatomical and cell cycle-dependent transcriptional shifts during heart development*

We pulled the UMI data uploaded by the authors at GEO (GSE122403). We subsequently performed UMAP + clustering through Seurat and selected clusters clearly identifiable as cardiomyocytes by marker gene expression.

*Sean Murphy et al. (Chulan Kwon)*

*Single-Cell Analysis Identifies PGC1 as a Master Regulator of Cardiomyocyte Maturation*

This data was generated in our lab; the raw data is currently not publicly available, but will be shortly. We selected only the wild-type cardiomyocytes for further analysis. Both 3’ counts and UMIs (output from zUMIs, using intronic and exonic reads) were used for analysis.

*Seitaro Nomura and Masahiro Satoh et al. (Hiroyuki Aburatani, Issei Komuro)*

*Cardiomyocyte gene programs encoding morphological and functional signatures in cardiac hypertrophy and failure*

The uploaded data of the authors does not contain mitochondrial reads. Therefore, we pulled and mapped the FastQ data from the sham cardiomyocytes from ENA (PRJNA376183) using STAR/FeatureCounts. However, we found that the dataset had a high percentage of mitochondrial reads; thus, we discarded this dataset.

*Blanca Pijuan-Sala, Jonathan Griffiths, and Caroline Guibentif et al. (John Marioni and Berthold Gottgens)*

*A single-cell molecular map of mouse gastrulation and early organogenesis*

We pulled the UMI data following the instructions from the authors’ Github page (https://github.com/MarioniLab/EmbryoTimecourse2018/blob/master/download/download.sh). We subsequently used the authors’ labelings to select cells classified as “Cardiomyocyte.”

*Zongna Ren and Peng Yu et al. (Li Wang)*

*Single-Cell Reconstruction of Progression Trajectory Reveals Intervention Principles in Pathological Cardiac Hypertrophy*

We pulled the UMI data uploaded by the authors at GEO (GSE120064). We subsequently performed TSNE + clustering through Seurat and selected clusters clearly identifiable as cardiomyocytes by marker gene expression. Many clusters had an abnormally high percentage of mitochondrial reads (e.g. *>*65%), so we selected only clusters with consistently reasonable mitochondrial percentages. We subsequently included only the control (non-disease) cells in our final analysis.

*Konstantina-Ioanna Sereti, Ngoc Nguyen, and Paniz Kamran et al. (Reza Ardehali)*

*Analysis of cardiomyocyte clonal expansion during mouse heart development and injury*

The uploaded data of the authors does not contain mitochondrial reads. Therefore, we pulled the FastQ data from ENA (PRJNA427266) and remapped using Kallisto (pseudo in batch mode with the -quant flag). We found the p1 timepoint to be bimodal in terms of mitochondrial gene expression. However, because of the relatively small cell number and large fraction of poor quality cells, we found that our top5 filter did not catch all low quality cells. We thus also excluded all p1 cells with >30% mitochondrial reads.

*Tabula Muris Consortium*

*A Single Cell Transcriptomic Atlas Characterizes Aging Tissues in the Mouse*

We pulled the BAM files for the Fluidigm studies from the publicly available AWS bucket (https://registry.opendata.aws/tabula-muris-senis/) and the annotations from the authors’ Figshare (https://figshare.com/projects/Tabula_Muris_Senis/64982). Specifically, for the latter, we used the metadata stored in the scanpy object, and matched the names of cells between this annotation and the raw data available through AWS. We focused only on the Fluidigm data for our study. We subsequently selected all cells from the “heart” tissue category. As we were unsure of the settings used to count with HTSeq, we recounted from the BAM files using FeatureCounts.

*Yin Wang and Fang Yao et al. (Li Wang)*

*Single-cell analysis of murine fibroblasts identifies neonatal to adult switching that regulates cardiomyocyte maturation*

We pulled the UMI uploaded by the authors at GEO (GSE122706). We subsequently performed TSNE + clustering through Seurat and selected clusters clearly identifiable as cardiomyocytes. However, we observed that the p14 cells demonstrates particularly high mitochondrial percentages, likely due to use of non-perfusion-based dissociation at this relatively late stage. We therefore discarded this timepoint.

*Florian Wunnemann (Gregor Andelfinger)*

*Currently unpublished cardiac dataset*

Counts table and metadata were kind provided by the authors. We used UMAP + clustering through Seurat to select cardiomyocytes based on clusters expressing cardiomyocyte markers and having comparable read and gene counts to other clusters. We additionally annotated clusters as atrial or ventricular, though we analyzed both sets of cells. However, as with several other Drop-seq datasets, we found that the data had unusually high entropy (and perhaps owing to low depth); thus, we discarded this dataset.

*Yang Xiao et al. (James Martin)*

*Hippo Signaling Plays an Essential Role in Cell State Transitions during Cardiac Fibroblast Development*

We pulled the UMI data uploaded by the authors are GEO (GSE100861). The metadata was kindly provided by the authors, and included the clustering used in the manuscript. We selected wild-type cells in the cardiomyocyte cluster. However, as with several other Drop-seq datasets, we found that the data had unusually high entropy (and perhaps owing to low depth); thus, we discarded this dataset.

*Haiqing Xiong, Yingjie Luo, Yanzhu Yue, and Jiejie Zhang et al. (Albin He)*

*Single-Cell Transcriptomics Reveals Chemotaxis-Mediated Intraorgan Crosstalk During Cardiogenesis*

The uploaded data of the authors does not contain mitochondrial reads. Therefore, we pulled and mapped the FastQ data from the four heart samples from ENA (PRJNA429249) using the FASTerQ approach with Kallisto|Bustools. Read 1 was used as the cdna read (base pairs 45 to 140, to avoid potential adaptor and poly A tails), while Read 2 was used for barcode (first 8 bp) and UMI (second 8 bp). Because of the short UMI length, we found that Kallisto|Bustools discarded many UMIs during counting. We thus used a custom script to count UMIs from the Kallisto|Bustools output.

*Michail Yekelchyk et al. (Thomas Braun)*

*Mono- and multi-nucleated ventricular cardiomyocytes constitute a transcriptionally homogenous cell population*

We pulled the mapped BAM files of wild-type cells from ENA (PRJEB29049) and recounted using FeatureCounts. However, we found that the dataset had high mitochondrial read percentage, and thus we discarded this dataset.

### Human In Vivo

*Michaela Asp and Stefania Giacomello et al. (Joakim Lundberg)*

*A Spatiotemporal Organ-Wide Gene Expression and Cell Atlas of the Developing Human Heart*

We pulled the UMI data and annotations for the single cell sequencing data from the authors’ website (https://www.spatialresearch.org/resources-published-datasets/doi-10-1016-j-cell-2019-11-025/). We used the authors’ clustering and selected all cells classified as cardiomyocytes.

*Yueli Cui, Yuxuan Zheng, and Xixi Liu et al. (Jie Qiao, Fuchou Tang)*

*Single-Cell Transcriptome Analysis Maps the Developmental Track of the Human Heart*

The uploaded data of the authors does not contain mitochondrial reads. Therefore, we pulled and mapped the FastQ data from the four heart samples from ENA (PRJNA415637) using the FASTerQ approach with Kallisto|Bustools. Read 1 was used as the cdna read (base pairs 45 to 140, to avoid potential adaptor and poly A tails), while Read 2 was used for barcode (first 8 bp) and UMI (second 8 bp). We then performed UMAP + clustering in Seurat, and selected clusters that clearly expressed cardiomyocyte markers. Because of the short UMI length, we found that Kallisto|Bustools discarded many UMIs during counting. We thus used a custom script to count UMIs from the Kallisto|Bustools output. We subsequently found that some, though not all, samples had unusually high mitochondrial percentages (namely - HE13W RV; HE17W AV, LA, LV, TV; HE20W RA; HE25W all samples). We also removed samples as post-filtering, there were too few for useful analysis (namely - HE23W, HE24W).

*Makoto Sahara and Federica Santoro et al. (Kenneth Chien)*

*Population and Single-Cell Analysis of Human Cardiogenesis Reveals Unique LGR5 Ventricular Progenitors in Embryonic Outflow Tract*

We downloaded the raw FastQ data from ENA (PRJNA510181), and subsequently mapped using Kallisto (pseudo in batch mode with the -quant flag). We had some concerns about several timepoints in this study due to high mitochondrial percentage. We eliminated some (namely - HE7W OFT, A; and HE8W); however, we are somewhat unsure about the quality of the data HE7W onwards.

*Hemant Suryawanshi et al. (Jill Buyon, Thomas Tuschl)*

*Cell atlas of the foetal human heart and implications for autoimmune-mediated congenital heart block*

The UMI data for the wild-type hearts was kindly provided by the authors as a Seurat object, and also included the authors’ UMAP clustering. We utilized their clustering to identify and select cardiomyocytes; we additionally filtered out cells with notably high mitochondrial percentage or low counts/genes.

*Li Wang, Peng Yu, Bingying Zhou, and Jiangping Song et al. (Shengshou Hu)*

*Single-cell reconstruction of the adult human heart during heart failure and recovery reveals the cellular landscape underlying cardiac function*

We download the UMI data and phenotype tables provided by the authors at GEO (GSE109816), selecting only the healthy heart tissue data. We used the authors’ provided metadata to select cardiomyocytes. We found that three of the four ventricular donors had extremely high mitochondrial percentages (∼70%). Very little is currently known about human adult CMs, so it is difficult to assess the validity of this range. However, one ventricular sample had a lower percentage, which also matched the atrial samples. We chose to therefore exclude the three samples with extremely high mitochondrial percentage, pending discovery of further information.

### Human Directed Differentiation

*Sherri Biendarra-Tiegs et al. (Timothy Nelson)*

*Single-Cell RNA-Sequencing and Optical Electrophysiology of Human Induced Pluripotent Stem Cell-Derived Cardiomyocytes Reveal Discordance Between Cardiac Subtype-Associated Gene Expression Patterns and Electrophysiological Phenotypes*

The counts data was kindly provided by the authors. We selected cardiomyocytes using the annotations provided in Figure 4 of the manuscript.

*Jared Churko et al. (Nathan Salomonis, Joseph Wu)*

*Defining human cardiac transcription factor hierarchies using integrated single-cell heterogeneity analysis*

We pulled the UMI data for all timepoints from the authors’ Synapse (https://www.synapse.org/#!Synapse:syn18078447/files/), using the V2 chemistry.

*Clayton Friedman, Quan Nguyen, and Samuel Lukowski et al. (Joseph Powell, Nathan Palpant)*

*Single-Cell Transcriptomic Analysis of Cardiac Differentiation from Human PSCs Reveals HOPX-Dependent Cardiomyocyte Maturation*

We pulled the UMI data for all timepoints from the authors’ processed data upload at ArrayExpress (E-MTAB-6268, https://www.ebi.ac.uk/arrayexpress/experiments/E-MTAB-6268/samples/).

*Kathryn Gerbin, Tanya Grancharova, Rory Donovan-Maiye, and Melissa Hendershott et al. (Ruwanthi Gunawardane)*

*Cell states beyond transcriptomics: integrating structural organization and gene expression in hiPSC-derived cardiomyocytes* We pulled the UMI data for all timepoints from the authors from the authors’ data upload at Quilt (https://open.quiltdata.com/b/allencell/tree/aics/integrated_transcriptomics_structural_organization_hipsc). However, we found that Batch 2 (composed of D0/D93/D96 cells) was under our approximate depth threshold (with a potentially similar issue to our Drop-seq data); thus, we focused on Batch 1. We performed TSNE + clustering through Seurat to identify cardiomyocytes.

*Elisa Giacomelli, Viviana Meraviglia, and Giulia Campostrini et al. (Valeria Orlova, Milena Bellin and Christine Mummery) Human-iPSC-Derived Cardiac Stromal Cells Enhance Maturation in 3D Cardiac Microtissues and Reveal Non-cardiomyocyte Contributions to Heart Disease*

We pulled the UMI data uploaded by the authors at GEO (GSE147694). We subsequently performed TSNE + clustering in Seurat to identify cardiomyocytes. We included only the control CMs (e.g. not co-cultured with ECs) in our final analysis.

*Hang Ruan and Yingnan Liao et al. (Leng Han, Li Wang)*

*Single-cell reconstruction of differentiation trajectory reveals a critical role of ETS1 in human cardiac lineage commitment*

We pulled the UMI for the D9, D14, and D60 timepoints from the tables uploaded by the authors at GEO (GSE129987).

*Adam Selewa et al. (Sebastian Pott, Anindita Basu)*

*Systematic Comparison of High-throughput Single-Cell and Single-Nucleus Transcriptomes during Cardiomyocyte Differentiation* We pulled the UMI data for the Drop-seq samples from the tables uploaded by the authors at GEO (GSE129096). However, as with several other Drop-seq datasets, we found that the data had unusually high entropy (and perhaps owing to low depth); thus, we discarded this dataset.

*Ana Silva et al. (Todd McDevitt)*

*Developmental co-emergence of cardiac and gut tissues modeled by human iPSC-derived organoids*

The UMI counts data was kindly provided by the authors. We performed TSNE + clustering through Seurat to identify cardiomyocytes.

### Mouse Direct Reprogramming

*Nicole R. Stone and Casey A. Gifford et al. (Deepak Srivastava)*

*Context-Specific Transcription Factor Functions Regulate Epigenomic and Transcriptional Dynamics during Cardiac Reprogramming*

We pulled the UMI data for the 3’ study from tables uploaded by the authors at GEO (GSE131328). We subsequently performed TSNE + clustering through Seurat and compared the generated clusters to those in the manuscript to assign cells into the putative trajectory reprogramming groups (as done by the authors).

**Supplementry Table 1:** In Vino Datasets

|  | Isolation | Sequencing | Mapping | Datatype | Depth | Genes | Cells | Timepoints | DOI | Species |
| --- | --- | --- | --- | --- | --- | --- | --- | --- | --- | --- |
| 10x Chromium Demo | 10x Chromium V3 | 10x Chromium | CellRanger | UMIs | 29833.0 | 4872 | 178 | e18 | 10x website | mouse |
| Asp and Giacomello | 10x Chromium V2 | 10x Chromium | CellRanger | UMIs | 11501.5 | 2899 | 762 | HE7W | doi:10.1016/j.cell.2019.11.025 | human |
| Cui, Zheng, and Liu | Picked | 8bp-STRT-seq | Kallisto/Bustools | UMIs | 33988.5 | 3834.5 | 2224 | HE9W, HE7W, HE6W, HE5W, HE25W, HE24W, HE23W, HE22W, HE20W, HE17W, HE15W, HE13W, HE10W | doi:10.1016/j.celrep.2019.01.079 | human |
| De Soysa | 10x Chromium V2 | 10x Chromium | CellRanger | UMIs | 17636.5 | 4331 | 6836 | e7.75, e8.25, e9.25 | doi:10.1038/s41586-019-1414-x | mouse |
| DeLaughter and Bick | Fluidigm C1 | Smart-seq2 | STAR/FeatureCounts | Reads | 950178.7 | 5751 | 630 | p0, e18, e14, e9, p3, e11, p21, p7 | doi:10.1016/j.devcel.2016.10.001 | mouse |
| Dong | Picked | 8bp-STRT-seq | Kallisto/Bustools | UMIs | 12444.0 | 2079 | 381 | e9.5, e10.5, e11.5 | doi:10.1186/s13059-018-1416-2 | mouse |
| Duan 10x | 10x Chromium V2 | 10x Chromium | CellRanger | UMIs | 6705.0 | 2137 | 2287 | p0 | doi:10.1016/j.celrep.2019.05.079 | mouse |
| Duan Drop-seq | Drop-seq | Drop-seq | Drop-seq Tools | UMIs | 4100.0 | 1700 | 5147 | p0 | doi:10.1016/j.celrep.2019.05.079 | mouse |
| Dueck | FACS | TruSeq | Kallisto | Reads | 25385903.8 | 13382 | 18 | e14.5 | doi:10.1186/s13059-015-0683-4 | mouse |
| Gladka and Molenaar | FACS | CEL-seq | BWA-ALN | UMIs | 23175.0 | 266 | 987 | p56 | doi:10.1161/CIRCULATIONAHA.117.030742 | mouse |
| Goodyer AVN | 10x Chromium V2 | 10x Chromium | CellRanger | UMIs | 10691.5 | 2710 | 2844 | e16.5 | doi:10.1161/CIRCRESAHA.118.314578 | mouse |
| Goodyer PF | 10x Chromium V2 | 10x Chromium | CellRanger | UMIs | 8832.0 | 2386 | 7177 | e16.5 | doi:10.1161/CIRCRESAHA.118.314578 | mouse |
| Hill e10.5 | 10x Chromium V2 | 10x Chromium | CellRanger | UMIs | 19166.0 | 4204 | 8375 | e10.5 | doi:10.1242/dev.180398 | mouse |
| Hill e13.5 | 10x Chromium V2 | 10x Chromium | CellRanger | UMIs | 6677.0 | 2270 | 13169 | e13.5 | doi:10.1242/dev.180398 | mouse |
| Jia, Preussner, and Chen | Fluidigm C1 | Smart-seq2 | STAR | Reads | 2831162.6 | 7033 | 421 | e8.5, e7.5, e9.5 | doi:10.1038/s41467-018-07307-6 | mouse |
| Kannan Chamber | LP-FACS | SCRB-seq | zUMIs | UMIs | 51454.0 | 4181.5 | 316 | p84 | doi:10.1161/CIRCRESAHA.119.315493 | mouse |
| Kannan Chamber 3' Counts | LP-FACS | SCRB-seq | zUMIs | 3' Counts | 361421.0 | 4181.5 | 316 | p84 | doi:10.1161/CIRCRESAHA.119.315493 | mouse |
| Kannan Fixed | LP-FACS | SCRB-seq | zUMIs | UMIs | 47500.5 | 4094.5 | 72 | p84 | doi:10.1161/CIRCRESAHA.119.315493 | mouse |
| Kannan Fixed 3' Counts | LP-FACS | SCRB-seq | zUMIs | 3' Counts | 847261.0 | 4094.5 | 72 | p84 | doi:10.1161/CIRCRESAHA.119.315493 | mouse |
| Kannan Reference | LP-FACS | mcSCRB-seq | zUMIs | UMIs | 14801.0 | 3000 | 936 | e18, e14, p0, p4, p14, p22, p28, p35, p56, p18, p11, p8 | none | mouse |
| Kannan Reference 3' Counts | LP-FACS | mcSCRB-seq | zUMIs | 3' Counts | 39128.0 | 3000 | 936 | e18, e14, p0, p4, p14, p22, p28, p35, p56, p18, p11, p8 | none | mouse |
| Lescroart, Wang, and Lin | FACS | Smart-seq2 | GSNAP/HTSeq | Reads | 1281114.0 | 10257 | 807 | e7.25, e6.75, e7.5, e6.5 | doi:10.1126/science.aao4174 | mouse |
| Li 10x | 10x Chromium V2 | 10x Chromium | CellRanger | UMIs | 9225.5 | 2685 | 4536 | e10.5 | doi:10.1242/dev.173476 | mouse |
| Li and Xu | Fluidigm C1 | Smart-seq2 | STAR/HTSeq | Reads | 1754692.0 | 7838 | 1829 | e9.5, e10.5, e8.5 | doi:10.1016/j.devcel.2016.10.014 | mouse |
| Murphy | LP-FACS | SCRB-seq | zUMIs | UMIs | 47668.0 | 7044 | 653 | p0, p7, p28, p21, p14 | doi:10.1101/2020.02.06.937797 | mouse |
| Murphy 3' Counts | LP-FACS | SCRB-seq | zUMIs | 3' Counts | 279638.0 | 7044 | 653 | p0, p7, p28, p21, p14 | doi:10.1101/2020.02.06.937797 | mouse |
| Nomura and Satoh | Picked | Smart-seq2 | STAR/FeatureCounts | Reads | 1670256.0 | 6292 | 89 | p56 | doi:10.1038/s41467-018-06639-7 | mouse |
| Pijuan-Sala, Griffiths, and Guibentif | 10x Chromium V1 | 10x Chromium | CellRanger | UMIs | 14718.0 | 3466.5 | 1206 | e7.75, e8.0, e8.5, e8.25 | doi:10.1038/s41586-019-0933-9 | mouse |
| Ren and Yu | iCell8 | Unknown | STAR/FeatureCounts | UMIs | 42270.5 | 2364.5 | 682 | 8w, 0w, 2w, 11w, 5w | doi:10.1161/CIRCULATIONAHA.119.043053 | mouse |
| Sahara and Santoro | Picked | Smart-seq2 | Kallisto | Reads | 1252637.4 | 15625 | 675 | HE6.5W, HE7W, HE8W, HE7.5W, HE5W, HE8.5W, HE5.5W | doi:10.1016/j.devcel.2019.01.005 | human |
| Sereti, Nguyen, and Kamran | Fluidigm C1 | Smart-seq2 | Kallisto | Reads | 387744.7 | 5714 | 123 | e9.5, e12.5, p1 | doi:10.1038/s41467-018-02891-z | mouse |
| Suryawanshi | 10x Chromium V2 | 10x Chromium | Drop-seq Tools | UMIs | 2854.0 | 998 | 4918 | HE19W, HE22W | doi:10.1093/cvr/cvz257 | human |
| Tabula Muris Senis | Picked | Smart-seq2 | STAR/FeatureCounts | Reads | 2306773.4 | 2375 | 441 | p720, p84 | doi:10.1101/661728 | mouse |
| Wang and Yao | iCell8 | Unknown | STAR/FeatureCounts | UMIs | 46481.0 | 3510 | 2205 | p1, p4, p7, p14 | doi:10.1038/s41467-020-16204-w | mouse |
| Wang, Yu, Zhou, and Song | iCell8 | Unknown | STAR/FeatureCounts | UMIs | 55018.0 | 2634 | 5229 | HP21Y, HP42Y, HP46Y, HP48Y, HP45Y, HP52Y, HP51Y, HP33Y, HP43Y, HP36Y | doi:10.1038/s41556-019-0446-7 | human |
| Wunnemann | Drop-seq | Drop-seq | Drop-seq Tools | UMIs | 3390.0 | 1460 | 26111 | e14.5, e16.5, e18.5, p1, p4, p7 | currently unpublished | mouse |
| Xiao | Drop-seq | Drop-seq | Drop-seq Tools | UMIs | 1567.0 | 1018 | 3673 | e13.5, e14.5 | doi:10.1016/j.devcel.2018.03.019 | mouse |
| Xiong, Luo, Yue, and Zhang | Picked | 8bp-STRT-seq | Kallisto/Bustools | UMIs | 23780.5 | 4556 | 1056 | e9.25, e8.25, e8.75 | doi:10.1161/CIRCRESAHA.119.315243 | mouse |
| Yekelchyk | iCell8 | SMARTer 3' | STAR/FeatureCounts | 3' Counts | 310298.7 | 4425 | 709 | p168 | doi:10.1007/s00395-019-0744-z | mouse |

**Supplementry Table 2:** PSC-CM and iCM Datasets

|  | Isolation | Sequencing | Mapping | Datatype | Depth | Genes | Cells | Timepoints | Doi | Species |
| --- | --- | --- | --- | --- | --- | --- | --- | --- | --- | --- |
| Biendarra-Tiegs | Fluidigm C1 | Smart-seq2 | MAP-Rseq | Reads | 3823726.0 | 9912 | 45 | D12, D40 | doi:10.1089/scd.2019.0030 | human |
| Churko | 10x Chromium V2 | 10x Chromium | CellRanger | UMIs | 11868.5 | 2758 | 13594 | D0, D5, D14, D45 | doi:10.1038/s41467-018-07333-4 | human |
| Friedman, Nguyen, and Lukowski | 10x Chromium V1 | 10x Chromium | CellRanger | UMIs | 8930.0 | 2416 | 44019 | D0, D2, D5, D15, D30 | doi:10.1016/j.stem.2018.09.009 | human |
| Gerbin, Grancharova, Donovan-Maiye, and Hendershott | FACS | SPLiT-seq | Drop-seq Tools | UMIs | 5535.5 | 2567 | 17470 | D24, D14, D12, D26, D96, D93, D0 | doi:10.1101/2020.05.26.081083 | human |
| Giacomelli, Meraviglia, and Campostrini | 10x Chromium V2 | 10x Chromium | CellRanger | UMIs | 5897.5 | 2057 | 6472 | D21 | doi:10.1016/j.stem.2020.05.004 | human |
| Ruan and Liao | iCell8 | Unknown | STAR/FeatureCounts | UMIs | 18607.5 | 2019 | 3600 | D9, D14, D60 | doi:10.1186/s12915-019-0709-6 | human |
| Selewa | Drop-seq | Drop-seq | STAR/FeatureCounts | UMIs | 1613.0 | 1057 | 37302 | D0, D1, D3, D7, D15 | doi:10.1038/s41598-020-58327-6 | human |
| Silva | 10x Chromium V2 | 10x Chromium | CellRanger | UMIs | 5783.0 | 2084 | 791 | D100 | doi:10.1101/2020.04.30.071472 | human |
| Stone and Gifford | 10x Chromium V2 | 10x Chromium | CellRanger | UMIs | 11302.0 | 3111 | 30728 | DM1, D1, D2, D3, D7, D14 | doi:10.1016/j.stem.2019.06.012 | mouse |

